# Single-Cell Atlas of Transcription and Chromatin States Reveals Regulatory Programs in the Human Brain

**DOI:** 10.64898/2026.02.02.703166

**Authors:** Yang Xie, Lei Chang, Guojie Zhong, Jonathan A. Rink, Tatiana Báez-Becerra, Ethan Armand, Wubin Ding, Kai Li, Eric Bonne, Audrey Lie, Hannah S Indralingam, Keyi Dong, Timothy Loe, Bohan Huang, Zhaoning Wang, Ariana S. Barcoma, Jackson K. Willier, Kyle W. Knutson, Jiayi Liu, Silvia Cho, Stella Cao, Kaitlyn G. Russo, Carissa K. Young, Jessica Arzavala, Yareli Sanchez, Aleksandra Bikkina, Natalie Schenker-Ahmed, Colin Kern, Zoey Zhao, Amit Klein, Jesus Flores, Chu-Yi Tai, Jacqueline Olness, Alexander Monell, Siavash Moghadami, Cesar Barragan, Chumo Chen, William Owens, Carolyn O’Connor, Michelle Liem, Mikayla V. Marrin, Cynthia Rose, Shane N. Alt, Nora Emerson, Julia Osteen, Jacinta Lucero, Daofeng Li, Rebecca D. Hodge, Ting Wang, C. Dirk Keene, Xiangming Xu, Quan Zhu, Joseph R. Ecker, M. Margarita Behrens, Bing Ren

## Abstract

Directly measuring chromatin states alongside transcription is essential for understanding how cell-type-specific regulatory programs are established and maintained in the adult human brain. We present a large-scale single-cell multimodal atlas generated by jointly profiling transcriptome with active (H3K27ac) and repressive (H3K27me3) histone modifications across 18 brain regions. We profile >750,000 nuclei spanning 160 cell types and integrate these data with chromatin accessibility, DNA methylation, 3D genome architecture, and spatial transcriptome. This framework annotates >500,000 regulatory elements and resolves cell-type-specific chromatin states. We link enhancers to target genes, infer gene regulatory networks, and classify chromatin interactions, revealing neuron-enriched long-range Polycomb repression of developmental genes. Integrating these maps with GWAS data and sequence-based model prioritizes noncoding variants, effector genes, and vulnerable cell types for neuropsychiatric disorders. Finally, cross-species comparisons show conserved activation but more divergent repression. Together, this study provides a functional reference for interpreting noncoding variants, epigenetic memory, and brain organization.

**HIGHLIGHTS:**

- Joint single-cell profiling of transcriptomes with active or repressive histone modification in >750,000 nuclei across adult human brain.
- Chromatin state annotation of >500,000 candidate *cis*-regulatory elements distinguishes active enhancers from accessible and Polycomb-repressed regions.
- Cell-type-resolved regulatory networks and sequence-based deep learning model prioritize functional neuropsychiatric risk variants.
- Spatial epigenomic imputation reveals laminar layer-specific Polycomb repression programs.
- Integration with 3D genome architecture reveals neuron-specific super long-range chromatin loops silencing early developmental genes.
- Evolutionary analysis uncovers conserved active regulatory grammar but divergent repressive landscape.

## INTRODUCTION

The functional diversity of the human brain arises from highly coordinated gene regulatory programs that specify and maintain cellular identity across development and throughout the lifespan^1,2^. Although most neuronal and glial lineages are established during embryonic and fetal stages, their identities must be stably preserved for decades while retaining sufficient plasticity to support learning, adaptation, and experience-dependent remodeling^3^. The balance between stability and plasticity is achieved through epigenetic mechanisms that encode regulatory memory beyond DNA sequence. Among these, post-translational histone modifications play a central role by marking genomic regions for activation or repression, thereby regulating transcriptional potential in a cell-type- and context-dependent manner.

Two histone modifications are particularly informative for regulatory states. Acetylation of histone H3 at lysine 27 (H3K27ac) marks active enhancers, promoters and is closely coupled to transcriptional output, whereas trimethylation at the same residue (H3K27me3), deposited by Polycomb repressive complexes, enforces transcriptional silencing of alternative lineage programs and developmental regulators in mature cell types^4-6^. Together, these marks capture complementary regulatory logic: activation of cell-type-specific gene programs and durable repression of inappropriate or earlier developmental states. Dysregulation of these epigenetic processes has been increasingly linked to neurodevelopmental, neuropsychiatric, and neurodegenerative disorders, highlighting the importance of resolving chromatin regulation in the human brain^7-9^.

Over the past decade, large-scale single-cell transcriptomic studies have transformed our understanding of cellular diversity in brain by defining detailed taxonomies across development, adulthood, and diseases. Parallel efforts profiling chromatin accessibility and DNA methylation at single-cell resolution have also begun to illuminate the regulatory genome underlying this diversity^10-13^. However, these modalities provide only a partial view of regulatory states. Chromatin accessibility alone does not reliably distinguish active enhancers from poised or structurally permissive elements, nor does it provide information about Polycomb-mediated repression. DNA methylation offers insight into durable silencing but lacks the temporal and functional specificity needed to dissect dynamic regulatory programs^14,15^. Consequently, critical questions remain unresolved: which regulatory elements actively drive transcription in each cell type, which maintained repressed yet potentially inducible, and how these programs are organized within nuclear 3D genome architecture and across tissue space.

Interpreting noncoding genetic variants further underscores these limitations. Genome-wide association studies (GWAS) have identified thousands of risk variants for schizophrenia, bipolar disorder, autism spectrum disorder (ASD), and Alzheimer’s disease, the vast majority of which lie outside protein-coding sequence. Assigning these variants to relevant cell types, regulatory elements, and effector genes remains challenging without cell-type-resolved, functional annotation of chromatin state^16,17^. Moreover, repression is often treated implicitly as a passive absence of activity, despite growing evidence that Polycomb-mediated silencing imposes biologically meaningful constraints on gene regulation and cellular plasticity^5,18-20^.

Here, we addressed these questions by generating a comprehensive single-cell multimodal atlas of the adult human brain that jointly profiles transcription and histone modifications marking active or repressive chromatin. Using Droplet Paired-Tag, we simultaneously measured transcriptome together with H3K27ac or H3K27me3 in individual nuclei across 18 anatomically defined brain regions from three male donors. The resulting dataset comprises over 750,000 nuclei spanning 160 cell types, providing a unified framework that directly links transcriptional output to regulatory state. By Integrating these data with matched chromatin accessibility, DNA methylation, 3D chromatin interaction maps and spatial transcriptomics, we moved beyond catalogs of cCREs to define dynamic chromatin states with cell-type- and regional specificity, and to connect them to gene regulatory programs. Leveraging this dataset, we trained a deep-learning model to predict the cell-type-specific functional impact of noncoding variants. Together, this resource provides a functional reference for studying the regulatory architecture of the adult human brain, and a foundation for understanding how regulatory programs shape cell identity, spatial organization, evolution, and genetic risk for neurological diseases.

## RESULTS

### A Multimodal Single-Cell Atlas of Active and Repressive Chromatin States in the Adult Human Brain

To delineate the regulatory programs that maintain cellular identity in the adult human brain, we focused on two histone modifications that correspond to active and repressive chromatin states: H3K27ac, which marks active regions such as enhancers and promoters, and H3K27me3, which represents Polycomb-mediated repression^21-23^. We used Droplet Paired-Tag to jointly profile the nuclear transcriptome together with either H3K27ac or H3K27me3 in the same nucleus (Figure 1A)^24^. We sampled 18 anatomically defined brain regions from three neurotypical male donors, spanning the cerebral cortex (encompassing 15 regions including frontal, temporal, parietal, and occipital lobes), cerebellum (CB), and pons (PN) (Figure 1B and Table S1.1). This sampling strategy captured fine-grained cortical heterogeneity while also incorporating developmentally distinct populations in hindbrain.

**Figure 1.**
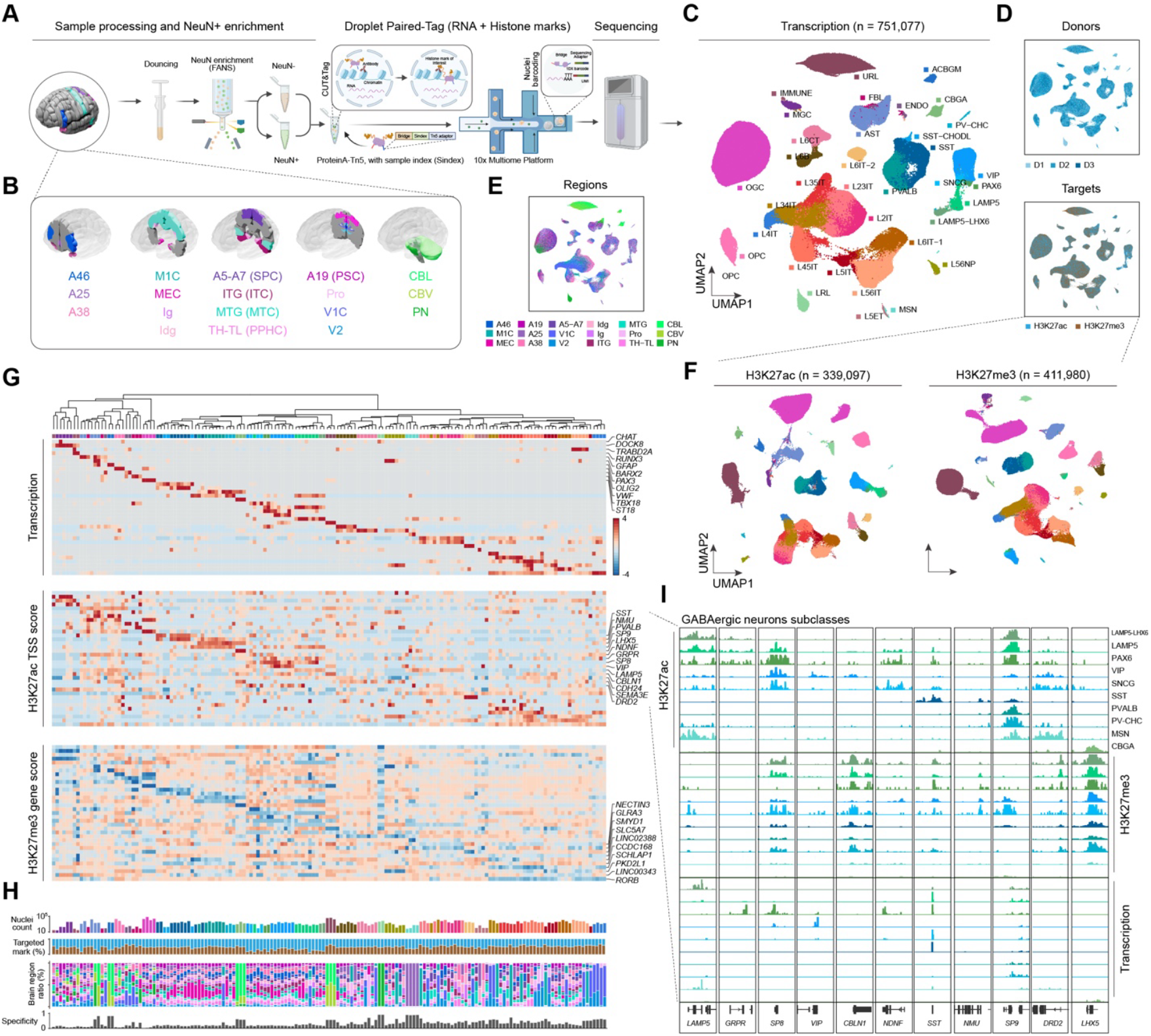
A single-cell joint histone modifications and transcriptomic atlas across 18 human brain regions. (A) Schematic overview of the experimental workflow, including sample processing, nuclei isolation, Droplet Paired-Tag single cell profiling and subsequent sequencing to simultaneously measure RNA and one of the two histone marks of interest (H3K27ac or H3K27me3) in single nucleus. (B) Anatomical map of the 18 sampled brain regions, grouped into three major structural categories: Cortex, Cerebellum (CBL, CBV), and Pons (PN) (C-E) Uniform Manifold Approximation and Projection (UMAP) visualization of the single-nucleus transcriptomic profiles (n = 751,077). Nuclei are colored by annotated cell subclass (C), donor identity and targeted histone mark (D), or brain region (E). (F) UMAP visualization of the single-nucleus H3K27ac (n = 339,097) and H3K27me3 (n = 411,980) profiles. Nuclei are colored based on transcriptomic cell subclass annotation. (G) Top: Hierarchical clustering of 160 cell clusters using transcriptomic profiles. Bottom: Heatmaps showing the pseudobulk RNA expression, H3K27ac signal at transcription starting site (TSS score), and the H3K27me3 signal at gene body (gene score) for representative marker genes across cell clusters. Values displayed have been Z-score normalized across cell types. Column annotation colors indicate cell subclass. (H) Summary bar plots displaying nuclei count (log scale, top), proportional distribution across targeted histone modifications and brain regions (middle), and the regional specificity score for each cell cluster (bottom). (I) Genome browser tracks of aggregated histone modifications and transcription signal for major GABAergic neuron subclasses at representative marker loci.

To improve neuronal representation and experimental throughput, we implemented two optimizations (Figure 1A). First, fluorescence-activated nuclei sorting (FANS) using NeuN staining was applied to balance the ratio of neuronal (NeuN+) vs non-neuronal (NeuN-) populations prior to single-cell profiling (Figure S1A and S1B). Second, we introduced short (2-3 bp) index sequence (hash indices) into the Tn5 transposase adaptors, enabling multiplexed processing of samples across regions or donors within a single droplet reaction (Table S1.2). These optimizations substantially increased the recovery of high-quality neuronal nuclei and reduced batch effects, with minimal influence on data complexity and signal enrichment (Figure S1C-J). Inspection of sequencing results also showed that data quality was comparable across donors at both library (Figure S1K-N) and single-nucleus levels (Figure S1O-Q).

After quality control and removal of doublets, the final dataset comprised 751,077 nuclei with paired transcriptomes and histone modification profiles, including 339,097 H3K27ac and 411,980 H3K27me3 profiles. Transcriptome-based clustering resolved three major cellular classes and 35 cell subclasses (Figure 1C, S2A and S3; Table S1.3). Across subclasses, non-neuronal nuclei contained fewer transcripts per nucleus but showed similar level of histone modification fragments per nucleus compared to neurons (Figure S2B). Notably, caudal ganglionic eminence (CGE) derived GABAergic interneurons exhibited fewer H3K27me3 fragments per nucleus than the other neuronal subclasses, a pattern not readily explained by sample preparation or quality metrics (Figure S2B). Despite this subclass-specific feature, transcriptomes robustly separated glutamatergic neurons, GABAergic neurons, and non-neuronal cells (Figure 1C). Nuclei from different donors, histone targets, and profiling strategies were well balanced across subclasses, indicating minimal technical bias (Figure 1D and S2C). As expected, glutamatergic and GABAergic neurons exhibited strong regional segregation, whereas non-neuron were more transcriptomically similar across regions (Figure 1E).

Projecting histone modification profiles onto the transcriptome-defined taxonomy demonstrated that histone modification data alone recapitulated the major cellular hierarchy observed in transcriptomes, while revealing additional regulatory resolution (Figure S2D and S2E). In particular, clustering analysis based on H3K27ac or H3K27me3 distinguishes forebrain and hindbrain populations within non-neuronal lineages such as oligodendrocytes (OGC) or astrocytes (AST) stronger than transcriptomes alone (Figure 1F and S3), suggesting that histone modifications capture developmental lineage-associated regulatory information that is not fully reflected by RNA expression.

Joint analysis of transcriptomes and histone profiles further resolved 160 distinct cell clusters, including rare and regionally restricted neuronal populations such as Purkinje cells in the cerebellum or underrepresented cortical interneuron subtypes (Figure 1G, 1H and S3; Table S1.4). These annotations agreed well with whole brain snRNA-seq and cortical SMART-seq references (Figure S4A-D)^8,9^. Across cluster-specific marker genes, H3K27ac at transcription start sites positively correlated with transcription level, while H3K27me3 over gene bodies inversely correlated with transcription and frequently marked lineage-restricted loci (e.g. *CHAT* and *TBX18* in GABAergic neurons) (Figure 1G). This regulatory “histone code” was particularly evident for lineage-defining genes across GABAergic subclasses (Figure 1I). For example, the transcription factor LHX5, a key Purkinje cells development regulator^25^, showed H3K27ac specifically in cerebellar GABAergic neurons (CBGA) and pervasive H3K27me3 in cortical interneurons, consistent with its RNA expression pattern (Figure 1I). Conversely, the *VIP* locus carried H3K27ac specifically in CGE-derived VIP interneurons. Although *VIP* expression was absent from medial ganglionic eminence (MGE) derived interneurons, this locus didn’t exhibit strong repression by H3K27me3 (Figure 1I).

Together, this dataset establishes a single-cell reference in which transcriptional identity is directly linked to active and repressive chromatin marks across the adult human brain, providing a foundation for functional annotation of regulatory elements and downstream integrative analyses.

### Chromatin State Annotation Resolves Functional Classes of Candidate Cis-Regulatory Elements in the Human Brain

Large-scale single-cell chromatin accessibility studies have generated an extensive catalog of candidate *cis*-regulatory elements (cCREs) in the human brain, providing a valuable resource for understanding regulatory genomics^26,27^. However, chromatin accessibility alone provides merely a permissive view of chromatin and does not distinguish between elements that are transcriptionally active, poised, silent or actively repressed. To resolve the functional states of cCREs while minimizing donor variation, we integrated our single-nucleus H3K27ac and H3K27me3 profiles with matched single-nucleus chromatin accessibility (CA) and DNA methylation (DNAme) data generated from the same donor cohort^26,28^.

Transcriptome was used as a common anchor to align histone modification, chromatin accessibility and DNA methylation modalities using canonical correlation analysis (CCA) (Figure 2A), yielding a unified cellular framework in which 32 transcriptomic subclasses were matched to 29 chromatin accessibility (CA) subclasses and 24 DNA methylation (DNAme) subclasses. (Figure S4E-P; Table S2.1). With each subclass, single-nucleus profiles from each modality were aggregated and binarized, and a multivariate Hidden Markov Model was trained using ChromHMM to infer genome-wide combinatorial chromatin states (Figure 2A)^29^. For DNA methylation, CpG hypermethylation scores were computed and binarized to accommodate its pervasive genomic distribution feature (STAR Method)^30^. This approach produced a nine-state model that captured the primary axes of regulatory variation and showed strong concordance with known genomic features or annotations (Figure 2B, S5A-D; Table S2.2)^31^.

**Figure 2.**
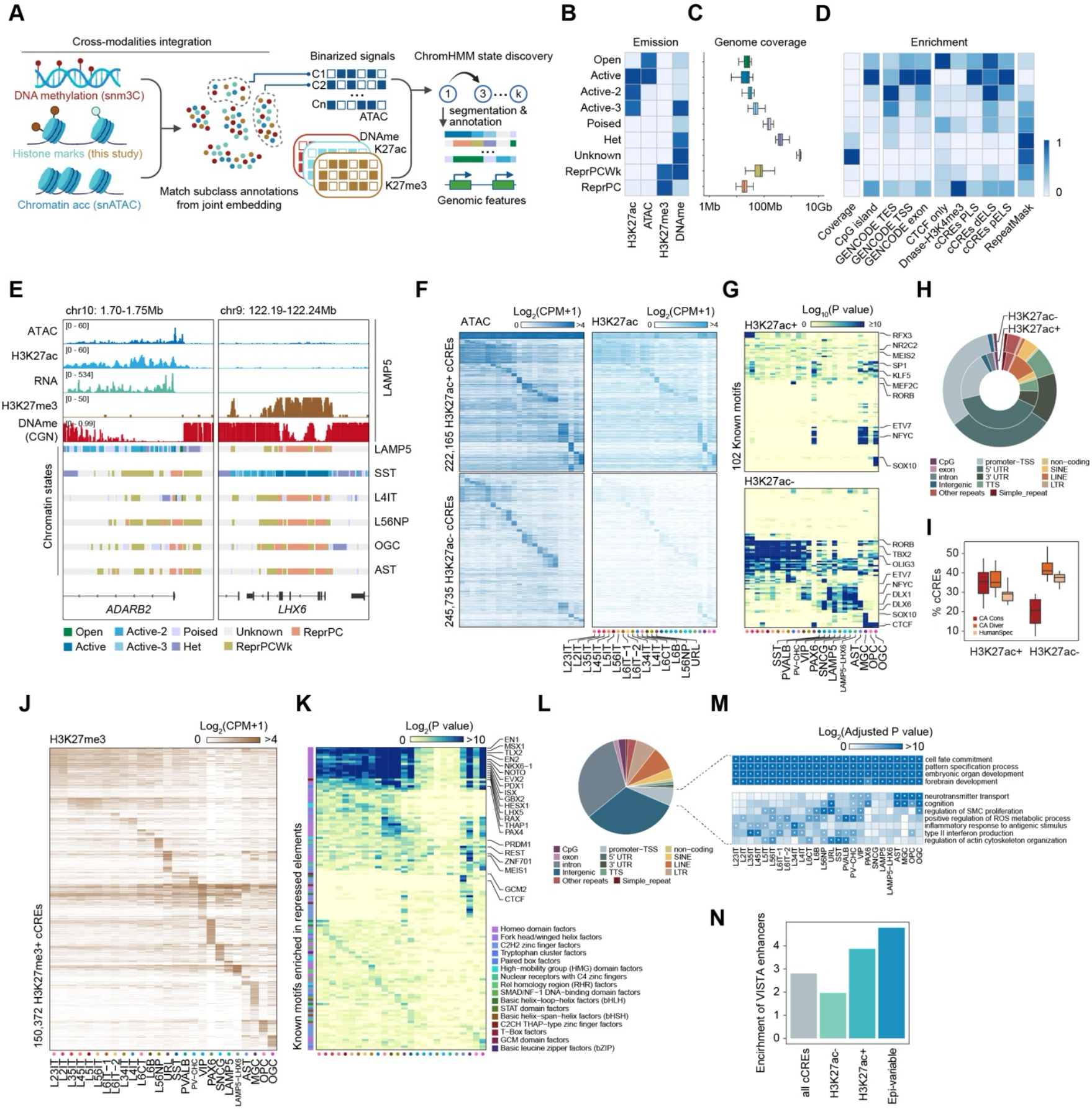
Annotation and characterization of candidate cis-regulatory elements (cCREs) with chromatin states. (A) Workflow for integrating DNA methylation (snm3C-seq), chromatin accessibility (snATAC-seq), and histone modifications to train a nine-state ChromHMM model. (B) Emission probabilities of the nine chromatin states for H3K27ac, ATAC, H3K27me3, and DNA methylation (DNAme). (C) Genome coverage of each chromatin state averaged over all cell subclasses (n = 32). For all boxes, the center line represents the median, box limits represent the upper and lower quartiles (75th and 25th percentiles), and whiskers represent 2× interquartile range (IQR). (D) Heatmaps showing the scaled genome coverage and overlapping genomic features including CpG island, GENCODE annotation, regulatory features from ENCODE SCREEN database (CTCF, Dnase-H3K4me3, cCREs PLS/dELS/pELS), and repetitive sequence (RepeatMask) for each chromatin state. Enrichment has been min-max normalized within each group. (E) Genome browser tracks showing aggregated epigenetic signals and resulting chromatin state segmentations at the *ADARB2* (CGE marker) and *LHX6* (MGE marker) loci for LAMP5 interneurons. Chromatin state segmentations for SST, L4IT, L56NP, OGC, and AST subclasses are also displayed. (F) Characterization of H3K27ac+ and H3K27ac-cCREs using chromatin states. Left: Heatmaps showing aggregated chromatin accessibility signal (log2 CPM+1) over H3K27ac+ and H3K27ac-cCREs across cell subclasses. Right: Heatmaps showing aggregated H3K27ac signal (log2 CPM+1) across the same cCREs and subclasses. Rows represent individual cCREs sorted by signal intensity. (G) Enrichment of known transcription factor motifs in H3K27ac+ and H3K27ac-cCREs across cell subclasses. Example motifs with distinct activities across subclasses or cCRE groups are labeled. Color indicates significance level (-log10 P value) calculated by hypergeometric tests in HOMER. (H) Genomic feature distribution of H3K27ac+ and H3K27ac-cCREs. (I) Box plot showing the percentage of cCREs classified as evolutionarily conserved (CA Cons) or divergent (CA Diver) or human specific (HumanSpec) based on chromatin accessibility conservation (Li *et al*., 2023) across subclasses. Box represents the interquartile range (IQR); whiskers extend to 2×IQR. (J) Heatmaps showing aggregated H3K27me3 signal (log2 CPM+1) over H3K27me3+ cCREs across cell subclasses. (K) Enrichment of known transcription factor motifs in H3K27me3+ cCREs across cell subclasses, similar as (G). Example motifs are labeled and colored based on families of their associated transcription factors. (L) Genomic feature distribution of H3K27me3+ cCREs, showing enrichment of repeat elements (e.g. SINE, LINE, LTR). (M) Gene Ontology (GO) enrichment analysis for genes associated with H3K27me3+ promoters. Ontology terms are segregated into development-restricted (top) versus lineage-restricted (bottom) functions. P values are calculated by hypergeometric test and adjusted using the Benjamini-Hochberg (BH) method. Asterisk indicates significant association (Adjusted P value < 0.05). (N) Bar plot showing the fold-enrichment of VISTA-validated enhancers within different cCRE classes relative to the genomic background. “Epi-variable” cCREs are defined as regions switching between H3K27ac+ and H3K27me3+ states across subclasses. Enrichment (odd ratio) is calculated by Fisher’s exact test.

Based on emission probability and genomic enrichment patterns (Figure 2B-D), the inferred chromatin states were grouped into four functional categories: open, active, Polycomb-repressed, and DNAme-enriched (Figure 2B). Importantly, incorporation of H3K27ac enabled separation of accessible elements that were active (“Active”, “Active-2,” and “Active-3” reflecting proximity to peak signal) from those lacking acetylation (“Open”). Active states were mostly enriched at promoters and enhancers, showing strong overlap with ENCODE-defined regulatory elements, whereas open states were preferentially enriched for structural features such as CTCF binding sites and showed weaker association with transcription (Figure 2D). Within the Polycomb-repressed category, the model distinguished promoter-proximal Polycomb-repressed regions with low DNA methylation (“ReprPC”; high H3K27me3, low DNAme) from a second state characterized by concurrent H3K27me3 and DNA methylation (“ReprPCWk”; high H3K27me3, high DNAme). The remaining states, including “Het” (Heterochromatin), “Poised” and “Unknown”, likely reflect constitutive heterochromatin, transitional regulatory configurations, unmarked regions or chromatin states marked by modifications not assayed here (Figure S5C).

Examination of individual loci across subclasses illustrated the specificity of chromatin state segmentations. For example, the CGE interneuron marker *ADARB2* adopted an active chromatin state (“Active”) in LAMP5 interneurons but switched to Polycomb-repressed state (“ReprPC”) in MGE interneurons as well as other subclasses (Figure 2E). Conversely, *LHX6*, a key MGE-interneuron marker, exhibited “ReprPC” state in LAMP5 interneurons, with strong H3K27me3 and DNAme across the gene body. These reciprocal patterns support a model in which cellular identity is encoded not only by activation of lineage markers but also by active repression of alternative fate programs.

Applying this framework to previously defined human brain cCREs enabled functional stratification of over 500,000 cCREs. After excluding six cell subclasses with insufficient coverage in at least one modality, on average 6.13% and 4.91% of cCREs per cell subclass were annotated as “Active” and “Open”, respectively, whereas nearly half (48.76%) were assigned to the “Unknown” state (Figure S5E). When restricting the analysis to accessible cCREs within each subclass, the fraction classified as “Active” and “Open” increased to 25.18% and 21.6%, respectively. We also identified a small fraction (2.08%) of Polycomb-associated accessible cCREs (Figure S5E). Consistent with functional relevance, “Active” cCREs showed the strongest enrichment for GWAS variants associated with neuropsychiatric disorders (e.g., schizophrenia), whereas “Open” state showed modest or no enrichment (Figure S5F), underscoring the value of state-resolved annotation for genetic interpretation.

Sequence and genomic feature analyses further separated “Active” from “Open” cCREs. To define sequence features associated with functional states, we compared H3K27ac+ (active) and H3K27ac-(open-only) cCREs across 26 subclasses (Figure 2F and S5G). The classification was supported by their distinct proximities to the nearest H3K27ac peak (Figure S5H). H3K27ac+ elements (n=222,165) were enriched for lineage-deterministic transcription factor (TF) motifs, including RORB in layer 4 intratelencephalic neurons (L4IT) and SOX10 in OGC (Figure 2G)^32,33^. In contrast, although H3K27ac-elements (n=245,735) also contained lineage-associated TF motifs (e.g. DLX1 and DLX6 in CGE interneurons)^34^, they frequently showed motif enrichments inconsistent with the resident lineage (e.g., SOX10 motifs in microglia), suggesting accessibility without lineage-appropriate activation. Relative to H3K27ac-elements, H3K27ac+ cCREs were more promoter-proximal (10.32% vs 1.04%), more often overlapped snm3C-seq loop anchors (59.36% vs 34.04%, Figure S5I), but were less enriched in repetitive elements such as LINEs (6.73% vs 14.70%, Figure 2H). H3K27ac+ sites also exhibited stronger evolutionary constraint in chromatin accessibility (33.12% vs 18.14%) and were more likely to have mouse-homologous sequences, whereas H3K27ac-sites were more human-specific (28.84% vs 38.00%; Figure 2I and S5J).

Repressive elements marked by H3K27me3 have remained under-characterized in the human brain, in part because fewer assays directly quantify repression at single-cell resolution. Using our chromatin state model, we identified 150,372 H3K27me3+ elements across 26 cell subclasses, with subclass-specific distributions that broadly mirrored those observed for H3K27ac (Figure 2J, S5K and S5L). Motif analysis revealed two distinct regulatory layers: a shared layer enriched for early developmental TFs, predominantly homeodomain TFs such as NKX6-1, EN2 and PDX1, and a subclass-specific layer with variable motif enrichment across subclasses (Figure 2K). Unlike active cCREs, H3K27me3+ elements were strongly enriched for repetitive sequences, including SINEs, LINEs, and LTR retrotransposons (Figure 2L). Analysis of genes with promoters classified as H3K27me3+ showed a similar two-tier pattern, combining broad repression of embryonic and developmental programs with lineage-restricted silencing in non-native contexts (Figure 2M). Together, these results support a model in which Polycomb-mediated repression contributes to both maintenance of cell identity and repression of transposable elements in terminally differentiated neurons.

Finally, to identify regulatory elements most critical for cellular identity, we defined 74,090 epigenetically variable (epi-variable) cCREs that switch between active (H3K27ac+) and repressed (H3K27me3+) states across subclasses. We hypothesized that these dynamic elements represent a functional core of the regulatory genome. To test this, we intersected different groups of cCRE with experimentally validated human enhancers from the VISTA Enhancer Browser^35^. Epi-variable cCREs showed the strongest enrichment, exceeding constitutively H3K27ac+ cCREs, the global cCRE background, and H3K27ac-cCREs (Figure 2N and S5M). These results suggest that elements undergoing active state transitions constitute a particularly potent and developmentally validated component of the human brain regulatory genome.

Overall, chromatin state annotation resolves functional heterogeneity among accessible regulatory elements, revealing how activation and Polycomb-mediated repression jointly shape cell-type-specific regulatory programs in the adult human brain.

### Cell-Type-Resolved Enhancer-Gene Links and Gene Regulatory Networks Connect Chromatin State to Transcription and Genetic Risk

A central challenge in regulatory genomics is assigning distal cCREs to their target genes in a cell-type-specific manner. To infer functional enhancer-gene relationships in the adult human brain, we integrated H3K27ac profiles with chromatin accessibility (snATAC-seq) and 3D chromatin interaction data (snm3C-seq) using the Activity-by-Contact (ABC) model (Figure 3A)^36^. This approach leverages enhancer activity (H3K27ac) and physical connectivity (3C data) to predict regulatory links and is well suited for histone-based measurement of activation.

**Figure 3.**
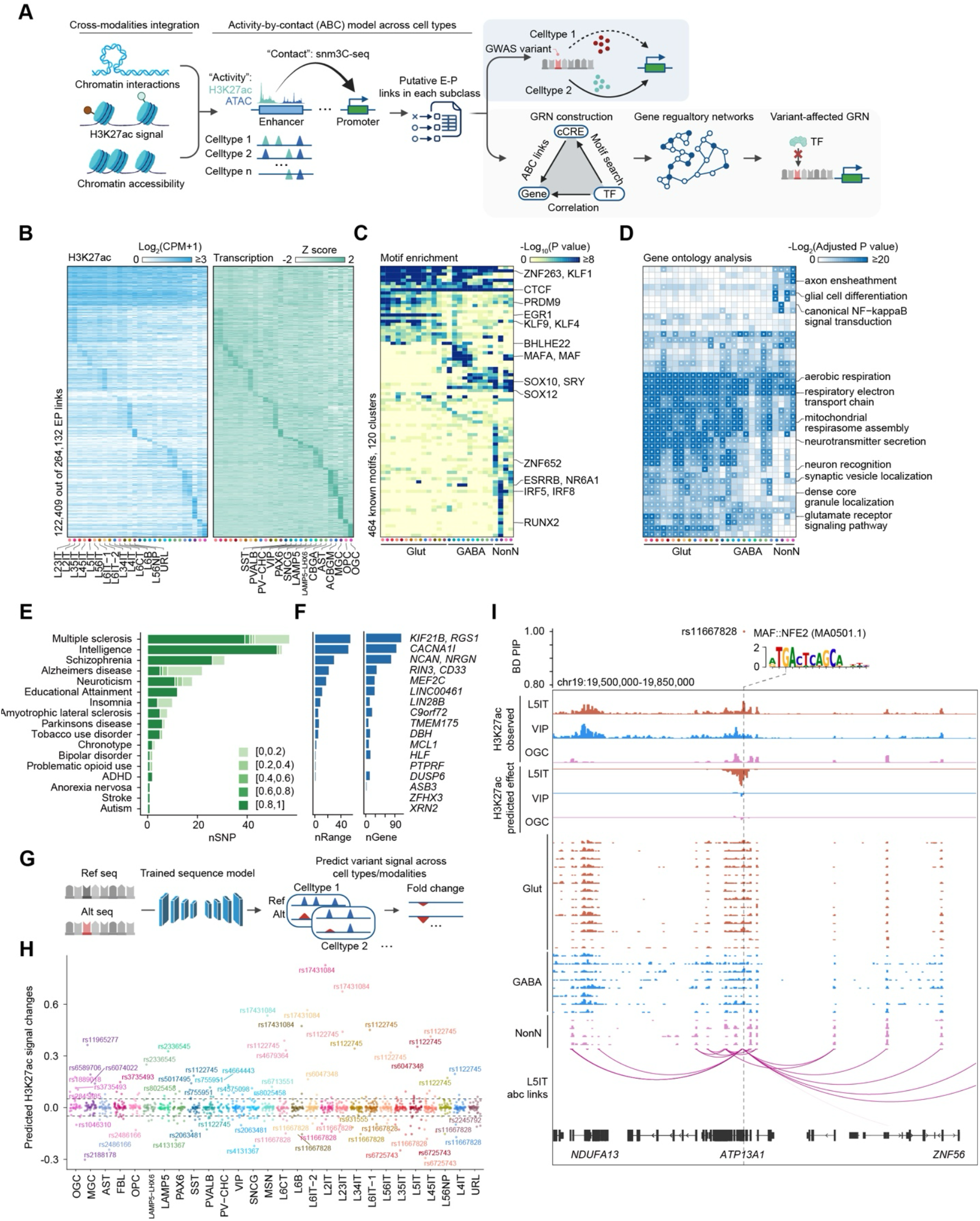
Genetic association of cell-type-specific regulatory links with neuropsychiatric diseases. (A) Schematic of the Activity-by-Contact (ABC) model used to link enhancers to genes. The predicted links were further used to prioritize variant effect and construct Gene Regulatory Networks (GRNs) for variant interpretation. (B) Heatmaps showing H3K27ac signal (log2 CPM+1) at predicted enhancers (left) and transcription of linked genes (RNA, right) across subclasses. RNA signals have been Z-score normalized. Rows are sorted using non-negative matrix factorization. (C) Enrichment of known transcription factor motifs in ABC links predicted enhancers across cell subclasses. Row represents motif clusters identified based on motif similarity. Example motifs in representative clusters with subclass-specific activities are labeled. Color indicates the highest significance level (-log10 P value) of motifs in each cluster, with P value calculated from hypergeometric tests calculated by HOMER. (D) GO analysis of target genes within each regulatory module. Color intensity represents -log2(Adjusted P value). P values are calculated by hypergeometric test and adjusted using the Benjamini-Hochberg (BH) method. Asterisk indicates significant association (Adjusted P value < 0.05). (E) Bar plot showing the number of fine-mapped variants (nSNP) for neuropsychiatric traits overlapping ABC-predicted enhancers. (F) Bar plots showing the total number of unique enhancers (nRange) and unique target genes (nGene) linked to risk variants for selected disorders. Example genes that known to be associated with the disease and linked to risk variants are displayed. (G) Schematic of the sequence-based deep learning model trained to predict variant effects on ATAC, H3K27ac, H3K27me3, and RNA signals. (H) Scatter plot of predicted H3K27ac signal changes (log scale) for fine-mapped variants across cell subclasses. Example variants with high predicted impact are labeled. (I) Characterization of cell type specific effects of a bipolar disorder fine-mapped variant (rs11667828). Top: Posterior inclusion probability (PIP) of all fine-mapped variants associated with BD in the displayed genomic range. MAF::NFE2 motif predicted to be affected by this variant is shown. Bottom: Genome browser tracks showing observed H3K27ac signal and model predicted variant effects on H3K27ac signal in representative cell types. ABC links in L5IT are also displayed.

Applying ABC model across cell subclasses yielded 332,713 predicted enhancer-gene links. After excluding four subclasses with insufficient chromatin interaction depth, we retained 264,132 high confidence links connecting 122,409 putative enhancers to 19,263 target genes across 28 cell subclasses (Figure 3B and S6A; Table S3.1). On average, each gene was linked to 2.66 enhancers and each enhancer to 1.90 genes, with a median enhancer-promoter distance of 84.7 kb (Figure S6B). Predicted enhancers and enhancer-gene pairs were significantly enriched for brain expression quantitative trait loci (eQTLs)^37-40^, including cell-type-specific eQTL-gene associations, supporting their functional relevance (Figure S6C-E).

Interrogating enhancer-gene links within each subclass further revealed a clear segregation of regulatory programs driven by distinct modules of TF motifs. For example, superficial layer intratelencephalic (IT) neurons were enriched for modules containing TFs such as EGR1 and PRDM9, linked to genes involved in glutamate receptor signaling and synaptic vesicle localization (Figure 3C and 3D). In contrast, oligodendrocyte (OGC) programs were dominated by SOX-family TFs (e.g., SOX10, SOX12) and modules enriched for genes required for axon ensheathment and glial differentiation (Figure 3C and 3D)^41-43^. These results demonstrate that enhancer-gene links anchored in chromatin activities and 3D genome architecture capture core features of cell-type-specific regulatory logic that underlies human brain cell identity.

To move from individual enhancer-gene links to a systematic overview of transcriptional regulation, we constructed gene regulatory networks (GRNs) by integrating enhancer-gene links with TF motif information and RNA expression. We adapted SCENIC+ to connect TFs to cCREs and downstream target genes (Figure 3A)^44^. Rather than relying on chromatin accessibility and pairwise correlations between cCRE activity and RNA expression, we used H3K27ac profiles together with ABC-predicted enhancer-gene links to build active GRNs. This approach identified 131 H3K27ac-associated GRNs, including 70 active GRNs in which TF expression was concordant with enhancer H3K27ac signal and target gene expression, and was significantly associated with at least one cell subclass (Figure S7A and S7B; Table S3.2).

These active GRNs recovered established lineage regulators across major brain cell types, including CUX1 and MBNL2 in glutamatergic neurons, DLX1 and LHX6 in GABAergic neurons, and RUNX1 in microglia, while also nominated candidate regulators (Figure S7B)^45-48^. We further used this framework to resolve regulatory differences among closely related interneuron lineages. Comparing MGE- and CGE-derived interneurons, we identified 17 active GRNs with lineage-biased activity (Figure S7C), recapitulating known determinants such as DLX1 and SP8 while nominating additional candidates including KLF5, ZBTB20 and ZNF704 (Figure S7D and S7E)^49^.

Incorporating H3K27me3 enabled the inference of repressive regulatory networks, which is largely inaccessible with accessibility-based approaches. We defined repressive GRNs as networks in which TF motif-associated cCREs carry H3K27me3 signal and inferred TF activity is inversely associated with target gene expression. Although most H3K27me3+ networks (59.2%) reflected a lack of activation based on their complementarity to active GRNs, we still identified 20 (16%) GRNs with a distinct repressive pattern, 17 of which were significantly associated with at least one cell subclass (Figure S7F; Table S3.3). We noticed that several TFs appeared in both active and repressive GRNs. For example, in glutamatergic neurons SATB2 is classically linked to activation of canonical targets, yet it showed strong negative association with GABAergic markers such as *GAD1* and *NXPH1*, which were marked by H3K27me3 in glutamatergic populations (Figure S7G and S7H). These results support a dual regulatory logic in which TFs can promote lineage programs while concurrently antagonizing alternative fate programs^50-52^.

The integration of enhancer-gene links and GRNs provides a solution for interpreting disease-associated noncoding variants. Using linkage disequilibrium score regression (LDSC), we partitioned SNP heritability from GWAS traits across H3K27ac+ cCREs and identified significant associations between 24 cell subclasses and 17 major neuropsychiatric traits (Figure S8A; Table S3.4). Among these traits, schizophrenia (SCZ) and bipolar disorder (BD) heritability were broadly enriched across neuronal subclasses, whereas Alzheimer’s disease (AD) heritability was preferentially enriched in microglia. To refine candidate variants, we performed statistical fine-mapping for major neuropsychiatric traits and intersected fine-mapped variants with the ABC-inferred regulatory links (STAR Method)^53,54^. This analysis identified 1,705 fine-mapped variants overlapping active enhancers within regulatory links across 30 GWAS traits, including 252 variants among 17 traits highlighted by LDSC (Figure 3E). The number of fine-mapped variants reflect each trait’s genetic architecture, GWAS sample size, and the underlying linkage disequilibrium structure^55,56^. Nonetheless, across major neuropsychiatric traits, we nominated a median of six candidate enhancers and 16 putative target genes per trait, with a median enhancer-gene distance of 19,291 bp for variant-overlapping enhancers (Figure 3F). Many nominated target genes aligned with prior genetic and functional evidence. For example, rs3852865 overlapped a microglial enhancer linked to *CD33*, a well-supported AD risk gene^57^. For BD, prior studies implicated *HLF*^58^, and our regulatory links identify a downstream enhancer (∼19 kb) overlapping rs9891134 in glutamatergic neurons as a candidate regulatory element for *HLF*.

To estimate variant effects at nucleotide resolution, we trained a sequence-based deep learning model that predicts cell-type-specific epigenetic signal and the allelic impact of sequence variants (Figure 3G; Also see Chang *et al*., co-submit). The model achieved high performance in predicting histone modification, chromatin accessibility and gene expression in held-out testing sequences across cell subclasses with sufficient depth (≥1000 nuclei) (Figure S8B). We applied the model to predict the cell-type-specific effect of H3K27ac signal fold-changes for all fine-mapped variants (posterior inclusion probability, PIP ≥ 0.1) and prioritized hundreds with large predicted effects on enhancer activity (Figure 3H and S8C; Table S3.5). One example is rs11667828, a fine-mapped BD risk variant. Although locates within the *ATP13A1* gene body, our regulatory links annotated this locus as a distal enhancer linked to *NDUFA13* and *ZNF56* (Figure 3I). The model predicted that the risk allele (G-to-T) disrupts a NFE2 motif and selectively reduces H3K27ac in layer 5 IT (L5IT) neurons. Consistent with this prediction, motif scanning supported disruption of a NFE2 binding site, and GRN analysis using default SCENIC+ settings implicated NFE2-related factors (e.g., NFE2L1) in regulating mitochondrial genes including *NDUFA13*, consistent with the proposed connections between mitochondrial dysfunction, reactive oxygen species (ROS) regulation, and BD susceptibility (Figure S8D)^59^.

Together, these analyses demonstrate how joint analysis of transcriptome and chromatin state enables the construction of cell-type-resolved regulatory maps to connect distal regulatory elements to target genes, define transcription factor networks, and translate noncoding GWAS associations into testable, cell-type-resolved regulatory mechanisms.

### Epigenetic Encoding of Structural and Areal Positional Identity in the Adult Human Brain

Anatomical compartmentalization of the human brain underlies distinct cognitive, sensory, and motor functions^60^, and is supported by region-specific cellular composition and gene regulatory programs^26,61^. While single-cell transcriptomic studies have revealed regional heterogeneity in neuronal populations, the extent to which epigenetic programs encode and maintain structural and areal identity remains unclear^9,62^. We therefore examined how histone modification landscapes vary across major brain structures and cortical areas within common cell subclasses.

We first assessed regional heterogeneity across the major brain structures profiled in this study: cerebral cortex, cerebellum, and pons. Because many neuronal subclasses are structure-restricted, we focused on non-neuronal populations to identify epigenetic determinants of structural identity. Differential gene expression analysis showed that astrocytes (AST) exhibit the most pronounced regional transcriptomic heterogeneity, whereas oligodendrocytes (OGC), oligodendrocyte precursor cells (OPC), and microglia (MGC) appear relatively homogeneous at the RNA level, consistent with prior reports (Figure 4A)^9,63^. In contrast, H3K27ac and H3K27me3 profiles revealed substantial regional heterogeneity in OGCs that was not apparent from transcriptomes alone (Figure 4A).

**Figure 4.**
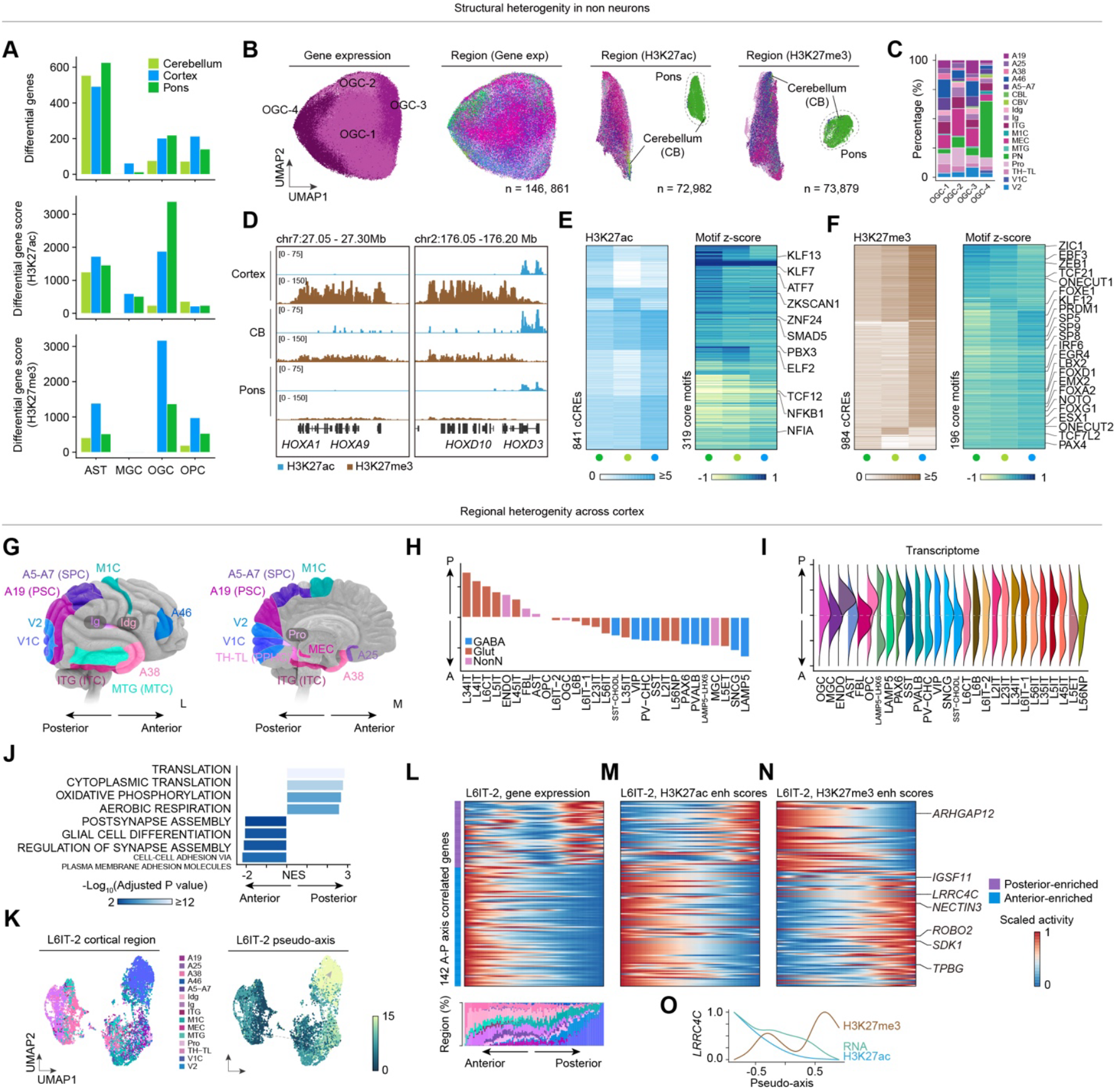
Structural and regional epigenetic variation in neural and glia populations. (A) Bar plots displaying the number of differentially expressed genes (top) and differential histone peaks (bottom) across three brain structures (Cortex, Cerebellum, Pons) for major glial subclasses (AST, MGC, OGC, OPC). (B) UMAP visualization of oligodendrocytes (OGC) based on transcriptomic (left), H3K27ac (middle), and H3K27me3 (right) profiles. Nuclei are colored by cell cluster annotations or brain region. (C) Stacked bar plot showing the regional composition of OGC cell clusters. (D) Genome browser tracks showing differential H3K27ac and H3K27me3 signals in OGC from different brain structures at *HOXA* and *HOXD* gene clusters. (E) Left: Heatmap of aggregated H3K27ac signal (log2 CPM+1) at regionally differential cCREs in OGC; Right: Average TF motif deviation scores at regionally differential motifs in OGC. Example motifs showing distinct activities are labeled. (F) Similar heatmaps as (E) but calculated with H3K27me3 signal. (G) Schematic of the cortical anterior-posterior (A-P) axis and regions selected for analysis. (H) Bar plot showing the Spearman’s rank correlation coefficients (SCC) of cell subclass abundance with the cortical A-P axis. Positive values indicate posterior enrichment; negative values indicate anterior enrichment. (I) Ridge plots of SCC quantifying the correlation between differential gene transcription signals and the A-P axis across subclasses. (J) Gene Set Enrichment Analysis (GSEA) of genes along the A-P axis in L6IT-2 neurons using correlation coefficients as rank. Color indicates significance level (-log10 Adjusted P value) calculated from fgsea using an adaptive multi-level split Monte-Carlo scheme and adjusted using the BH method. (K) UMAP visualization of L6IT-2 neurons. Nuclei are colored by brain region (left) or pseudo-axis score (right). (L-N) Top: Heatmaps showing the scaled transcription (L), H3K27ac enhancer scores (M), and H3K27me3 enhancer scores (N) for A-P axis correlated genes in L6IT-2 neurons, ordered by pseudo-depth. Value for each feature has been scaled between 0 and 1. Bottom: Bar chart showing the nuclei distribution across brain regions along the pseudo-axis bin. (O) Line plots showing the scaled activity of a representative anterior-enriched gene (*LRRC4C*) along the pseudo-axis for RNA, H3K27ac, and H3K27me3.

To quantify epigenetic heterogeneity, we estimated region-explained variance using PERMANOVA on low-dimensional embeddings (Figure S9A and STAR Methods)^64^. Across the four major non-neuronal populations, histone modification profiles captured markedly more region-associated variance (mean R^2^ = 0.29 for H3K27ac; 0.37 for H3K27me3) than chromatin accessibility (mean R^2^ = 0.09), a pattern further supported by the differences in silhouette score (Figure S9B). Consistent with these metrics, clustering analysis of histone modification profiles revealed robust regional separation across non-neuronal lineages including OGC (Figure 4B, 4C and S9C-E). In contrast, transcriptomic separation was modest for most non-neuronal subclasses except astrocytes (Figure S9E and S9F). These patterns suggest that oligodendrocytes retain region-specific regulatory programs encoded in histone modification landscapes despite relatively similar transcriptional outputs.

Inspection of specific gene loci highlighted the developmental origins of these structural signatures. Cortical OGCs showed strong H3K27me3 enrichment over *HOX* gene clusters (*HOXA, HOXC, HOXD*) or at loci such as *EMX1* and *FEZF2*, genes that are active in cortical progenitors and glutamatergic neurons. These repression patterns were largely absent in hindbrain (pons and cerebellar) OGCs, consistent with the established roles for these factors in axial patterning during development (Figure 4D)^65,66^. Beyond these loci, we identified 841 and 984 cCREs with regionally differential H3K27ac and H3K27me3 signals in OGCs, respectively. Motif analyses of histone modification signal further highlighted TF programs associated with structural identity. Cortical OGCs showed elevated activity of OGC lineage TFs such as TCF12 and NFIA, whereas hindbrain OGCs showed elevated activity of ELF2, KLF13, and KLF7 (Figure 4E)^67-69^. Conversely, ZIC1 motif was preferentially repressed in hindbrain OGCs, while NOTO and ONECUT2 motifs were more repressed in cortical OGCs (Figure 4F)^70,71^. Together, these findings indicate that structural identity in glial lineages is maintained through coordinated activation and Polycomb-associated repression of region-specific regulatory programs.

Astrocytes exhibited a similar but more pronounced form of structural heterogeneity. Both transcriptomic and epigenetic analyses separated telencephalic (cortical; ASCT) astrocytes from the non-telencephalic (cerebellum/pons; ASCNT) populations (Figure S9E-G). Relative to ASCNT, ASCT were characterized by strong transcriptomic and epigenetic activity of TFs such as FOXG1 and OTX1, both key regulators of forebrain development that were repressed in hindbrain-derived ASCNT (Figure S9H and S9I)^72-74^. These results further support a model in which epigenetic regulation preserves positional identity across diverse glial lineages in the adult human brain.

We next examined regional heterogeneity within the cerebral cortex, which is organized along a continuous anterior-posterior (A-P) functional axis rather than showing discrete structural boundaries^75-77^. We hypothesized that this graded organization is encoded in the epigenome of common subclasses. To test this, we quantified regional variation across 14 cortical regions for all subclasses using Local Inverse Simpson’s Index (LISI) (Figure 4G and S10A)^78^. Glutamatergic neurons exhibited the strongest regional heterogeneity in both transcriptomic and histone modification profiles, with the largest effects in L34IT and L4IT neurons, which represent specialized excitatory neuron populations enriched in visual cortex (V1C and V2) (Figure S10B). GABAergic neurons displayed weaker regional variation, and non-neuronal subclasses showed the least. Overall, cortical regional patterning was more pronounced in transcriptomes than in histone modification profiles (Figure S10A).

Consistent with these patterns, glutamatergic neurons showed the largest numbers of regionally differential genes and cCREs (Figure S10C, STAR Methods). Many of these features implicated core neuronal processes including ion transport. For example, in L35IT neurons, *CRH* and *CNR1* show higher RNA expression and enhancer activities in anterior regions, whereas *SCN1B, TRPC3*, and *HTR1B* were enriched posteriorly (Figure S10E)^9,79,80^. To determine whether these differences aligned with the continuous A-P axis, we first examined cell-type composition along the axis. Glutamatergic subclasses such as L6CT, L34IT, and L4IT were enriched in posterior regions, while GABAergic subclasses such as LAMP5 and SNCG were biased toward anterior regions. In comparison, non-neuronal populations showed weaker compositional bias (Figure 4H). We then identified axis-dependent features whose activity correlated with A-P position, nominating 3,438 genes and 12,379 (H3K27ac) and 5,569 (H3K27me3) axis-associated cCREs using an absolute correlation cutoff of 0.3 (Figure 4I and S10F, STAR Method). Among subclasses, L34IT and L6IT-2 showed particularly high number of posterior-biased cCREs (Figure S10F). Gene set enrichment analysis (GSEA) further revealed functional stratification along the axis. In L6IT-2, anterior cells were enriched for biological processes “synapse assembly” and “cell-cell adhesion”, whereas posterior L6IT-2 cells were enriched for “oxidative phosphorylation” and “translation”, consistent with distinct functions and metabolic demands across cortical areas (Figure 4J)^81,82^. To resolve these gradients at higher resolution, we applied diffusion maps on single-nucleus transcriptome profiles to derive a within-subclass pseudo-axis score, with L6IT-2 among the subclasses showing the clearest A-P gradient (Figure 4K)^83^.

Importantly, this transcriptional gradient was mirrored in the epigenome. For genes tightly coupled to the pseudo-axis, including *IGSF11* and *ROBO2*, we observed ABC-derived enhancer H3K27ac scores changed concordantly with RNA expression, and enhancer H3K27me3 scores shifted inversely (Figure 4L-4N). Notably, along the pseudo-axis, increase in enhancer H3K27ac scores preceded maximal increases in transcription, as exemplified at gene *LRRC4C* (Figure 4O). These coordinated epigenetic and transcriptional gradients provide evidence that continuous cortical areal identity is closely associated with graded regulatory states.

In summary, these analyses demonstrate that histone modification landscapes encode both discrete structural identities across brain structures and continuous areal specialization within cortex. By dissecting positional regulatory programs that are incompletely captured by transcriptomes alone, this resource provides a framework for understanding how spatial organization and developmental history are recorded and maintained in the adult human brain.

### Spatial Integration Reveals Laminar Epigenetic Programs in the Human Cerebral Cortex

The functional architecture of the human cerebral cortex is organized across laminar layers that segregate sensory inputs, intracortical computations, and output pathways^84,85^. This cytoarchitecture arises from the coordinated developmental programs that specify neuronal lineages and guide their migration to layer-specific destinations through distinct routes and timing^60,86,87^. These lineage histories and final laminar positions impose regulatory constraints, demanding precise epigenetic control to maintain layer-specific connectivity and physiological properties throughout life^88-92^. However, dissociation-based single-nucleus assays erase spatial context, limiting our ability to reconstruct the spatial logic that shapes gene regulation in the human cortex.

To restore spatial context to the epigenomic atlas, we performed Multiplexed Error-Robust Fluorescence in situ Hybridization (MERFISH)^93^ on tissues from four representative cortical regions across four donors distinct from those used for Droplet Paired-Tag. Using the MERSCOPE Ultra platform, we imaged tissues from all donors in parallel to minimize batch effects and generate highly consistent spatial transcriptomic profiles across replicates (Figure S11A-C). After stringent quality control, we recovered 362,900 nuclei across 20 slides and annotated them by referencing Droplet Paired-Tag profiles from matched dissections (Figure S12A-C; STAR Methods). The resulting spatial map recapitulated the expected segregation of neurons and glia and canonical laminar distributions of major subclasses, particularly among glutamatergic neurons (Figure S12D and S12E). Consistent with prior spatial transcriptomic studies and neuroanatomical literature, MGE-derived interneurons (SST and PVALB) spanned superficial to deep layers, whereas CGE-derived interneurons were preferentially enriched in superficial layers (Figure S12E and S12F)^12,94-96^.

To jointly interpret spatial and epigenomic measurements, we leveraged shared transcriptomic signatures as an anchor and imputed pseudo-spatial coordinates for each Droplet Paired-Tag nucleus by mapping it to its nearest MERFISH neighbors in a CCA-integrated embedding (Figure 5A; STAR Methods). This strategy effectively projects histone modification profiles onto the imaging coordinates, enabling analysis of regional and laminar epigenetic heterogeneity that was inaccessible by dissociation-based assays. As validation, the imputed laminar distributions of Droplet Paired-Tag nuclei closely matched the physical coordinates and marker gene activity in MERFISH data (Figure 5B, S12G and S12H), establishing a robust framework for high-resolution analysis of spatially structured epigenetic variation.

**Figure 5.**
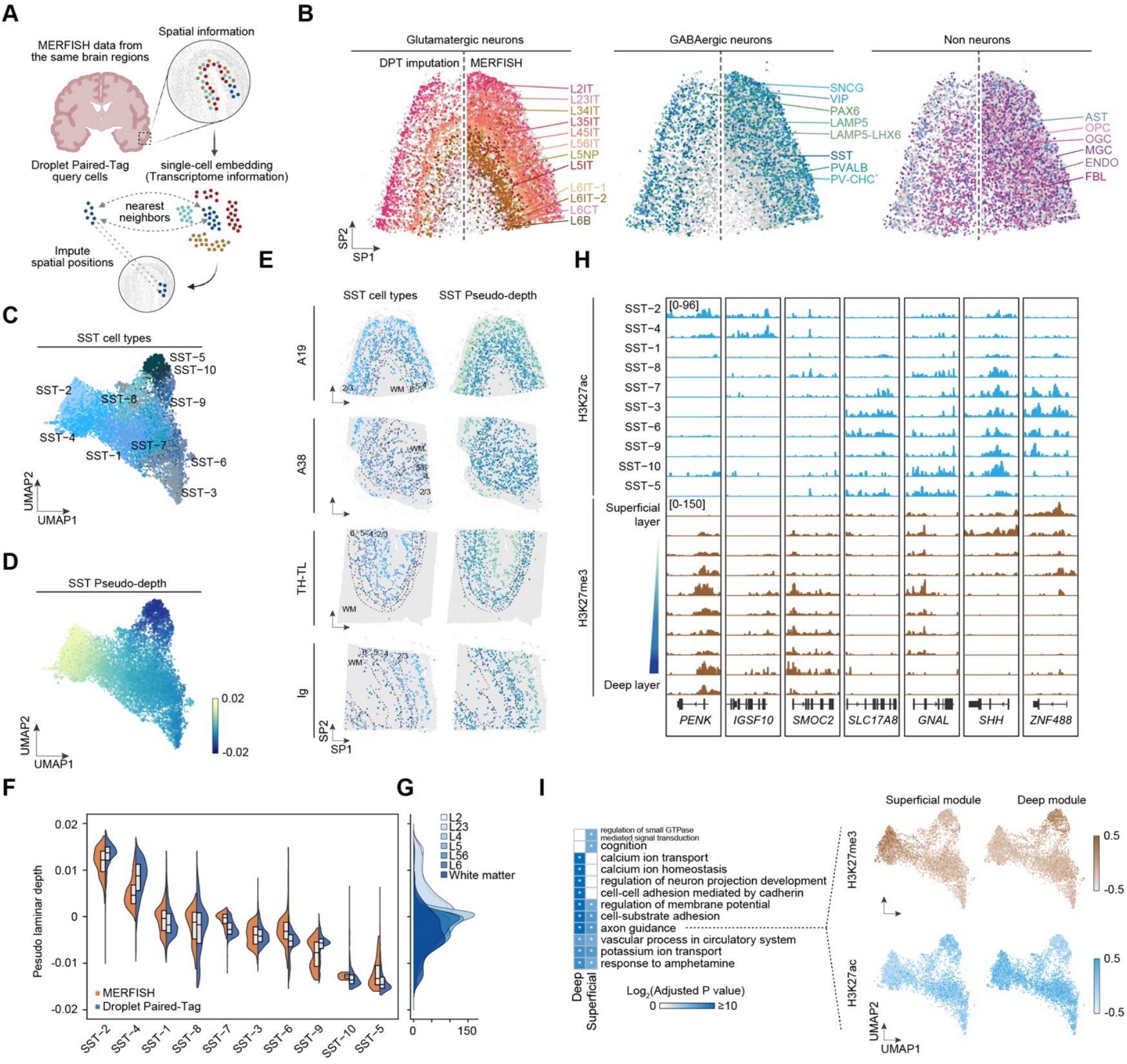
Spatial epigenomic integration revealed laminar identity of cortical interneurons. (A) Schematic of the multimodal integration strategy mapping Droplet Paired-Tag profiles to MERFISH spatial transcriptomics data to impute spatial coordinates. (B) Example spatial plot of MERFISH subclass annotation (right) or imputed spatial coordinates (left) for Droplet Paired-Tag nuclei for glutamatergic, GABAergic, and non-neuronal cells, colored by cell subclass. Slide is from brain region A19. (C-D) UMAP visualization of SST interneuron on the integrated embeddings, colored by SST cell clusters (C) or imputed pseudo-depth (D). (E) Spatial distribution of predicted SST cell clusters (left) and pseudo-depth (right) across different brain slices in MERFISH data. Contours indicate laminar layer classification. (F) Distribution of predicted pseudo-depth within each SST cell type, compared between MERFISH data and Droplet Paired-Tag data. Box represents the interquartile range (IQR); whiskers extend to 2×IQR. (G) The distribution of laminar layer classification along pseudo-depth in MERFISH cells. Pseudo-depth value is shown in (F). (H) Genome browser tracks of aggregated H3K27ac and H3K27me3 signals in SST cell clusters at representative gene loci. Genes with differential histone modification signals across laminar layers were selected for display. (I) Left: Gene ontology analysis of genes with differential H3K27me3 signal in superficial layer versus deep layer SST cell clusters, defined by pseudo-depth. Right: UMAP visualization of the scaled module scores calculated using the histone modification signals for axon guidance genes preferentially repressed in superficial SST (superficial module) or deep layer SST (deep module). P values are calculated by hypergeometric test and adjusted using the Benjamini-Hochberg (BH) method. Asterisk indicates significant association (Adjusted P value < 0.05).

We applied this framework to dissect epigenetic heterogeneity within somatostatin-expressing (SST) interneurons. SST interneurons arise predominantly from the dorsal-posterior medial ganglionic eminence, and populations destined for different cortical layers migrate with distinct timing and routes^97-99^. However, the regulatory programs associated with these layer-specific trajectories remain incompletely defined. We annotated 10 SST clusters in the MERFISH dataset that exhibited distinct spatial distributions (Figure 5C). Cell clusters such as SST-2 and SST-4 were enriched in superficial layers (L1-3) and expressed markers including *CALB1, IGSF10*, and *MME*. In contrast, SST-6 and SST-9 were biased toward deep layers (L4-6) and showed higher expression of *DCC* and *TRHDE* (Figure S12I)^96,100^. Diffusion map analysis further revealed that the first diffusion component (DC1) strongly correlated with cortical depth, providing a quantitative “pseudo-depth” that positions individual nuclei along the laminar axis (Figure 5B and 5C; STAR Methods)^83^. This pseudo-depth metric was robust across datasets and cell subtypes (Figure 5D and 5E), enabling systematic analysis of layer-associated variation in chromatin state.

Across pseudo-depth, specific loci showed coordinated shifts in the balance between H3K27ac and H3K27me3. For example, *PENK* displayed strong H3K27ac in superficial SST neurons but prominent H3K27me3 in deep-layer SST, while deep-layer markers such as *SMOC2* exhibited the opposite polarity (Figure 5F). Beyond cell type marker genes, we detected laminar-structured repression of developmental regulators. Superficial SST neurons accumulated H3K27me3 at ventral morphogen genes including *SHH, WNT5A*, and *WNT8B*, whereas deep-layer SST neurons preferentially repressed *WNT7A* and *BMP6* (Figure S12J)^101-103^. These reciprocal repression patterns suggest that superficial and deep-layer SST populations retain distinct epigenetic memories of their origins or the signaling environments during specification and migration. Consistent with this divergence, genes preferentially repressed in superficial SST were enriched for biological processes such as “small GTPase-mediated signaling” and “cognition”, whereas deep-layer SST showed stronger repression of genes involved in “calcium ion transport and homeostasis”, implicating layer-associated differences in calcium dynamics and electrophysiological properties aligned with distinct circuit outputs (Figure 5G). Notably, although axon guidance programs were repressed in both compartments, repression involved largely non-overlapping gene modules with complementary patterns of H3K27ac and H3K27me3 across SST subtypes and pseudo-depth (Figure 5H).

In summary, these analyses reveal that laminar identity in the adult human cortex is supported by spatially organized gene regulatory programs that integrate transcriptional activation with Polycomb-mediated repression. By projecting epigenetic heterogeneity onto physical space, our resource illuminates how developmental history and positional information persist in the adult brain and provides a framework for linking chromatin state to cortical architecture.

### Functional Annotation of Chromatin Loops Reveals Subclass-Specific Active and Repressive 3D Genome Topologies

Three-dimensional genome folding facilitates communications between distal regulatory elements and their target genes over large genomic distances^104-106^, yet chromatin interaction data alone provides limited insight into the functional nature of these contacts. To determine how chromatin topology reflects regulatory state in the adult human brain, we refined and annotated chromatin loops from snm3C-seq by intersecting 10-kb loop anchors with subclass-matched H3K27ac and H3K27me3 peaks (Figure 6A; STAR Methods)^107^. This strategy classified 74,410 loops across 32 transcriptomic subclasses, including 65,790 active loops and 5,366 repressive loops (Figure 6B and S13A). Across neuronal and non-neuronal subclasses, active loops dominated the annotated interactome (median 90.38%). These loops were typically short-range (<300 kb), more than half of them (51.92%) involved promoter-anchored contacts in which promoter activity was concordant with the distal anchor state (Figure 6C and 6D).

**Figure 6.**
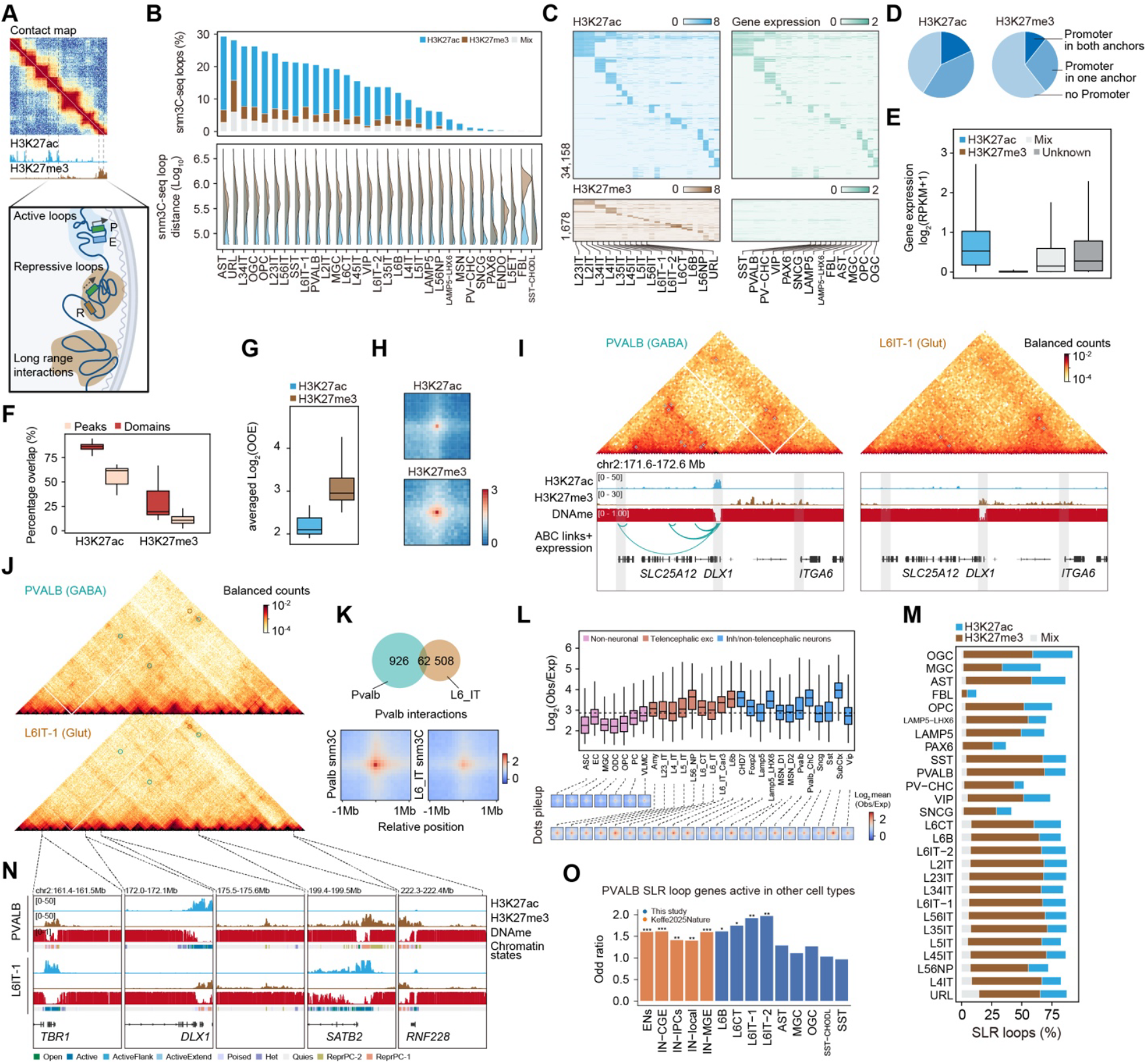
Annotation of subclass-specific short-range and super long-range snm3C-seq interactions. (A) Schematic of chromatin interactions classified as active, repressive, and super long-range (SLR) loops. (B) Top: Percentage of snm3C-seq loops classified as active, repressive, or “mixed” across cell subclasses. Bottom: Ridge plots showing the genomic distance distribution for active and repressive interactions across subclasses. (C) Top: Heatmap of H3K27ac signal (log2 CPM+1) at anchors of active loops associated with at least one promoter. Averaged transcription signals (log2 RPKM+1) of the associated genes with promoters in each anchor are displayed for comparison. Bottom: Heatmap of H3K27me3 signal (log scale CPM) at anchors of repressive loops associated with at least one promoter. Transcription signals are also displayed. (D) Pie charts showing the proportion of active and repressive loops involving promoters. (E) Box plots of gene expression (log2 RPKM+1) for genes anchored in active, repressive, or “mixed” loops. (F) Box plots showing the percentage overlap of active and repressive loop anchors with narrow peaks vs. broad domains. (G) Box plots of Aggregate Peak Analysis (APA) scores (log2 Observed/Expected contact ratio) for active and repressive loops. (H) APA heatmaps showing contact enrichment for active and repressive loops. (I) Top: Heatmap of balanced chromatin contacts in PVALB versus L6IT-1 at the *DLX1* locus at 10kb resolution. Bottom: Genome browser tracks showing the aggregated H3K27ac, H3K27me3 DNA methylation signals as well as ABC links in PVALB and L6IT-1 cell subclass. (J) Heatmap of balanced chromatin contacts in PVALB versus L6IT-1 at the *DLX1* locus at 100kb resolution. Subclass-specific super long-range chromatin interactions (> 5Mb) in PVALB versus L6IT-1 are highlighted. (K) Venn diagram of SLR interactions between PVALB and L6IT-1 (top), and pile-up APA heatmaps showing subclass-specific chromatin interaction signals at PVALB-specific SLR loops (bottom). (L) Top: Box plots of normalized SLR interaction strength across cell subclasses from Tian *et al*., 2023. Bottom: Pile-up heatmaps of snm3C-seq signal over the identified SLR loops across subclasses. (M) Percentage of SLR loops classified as active (H3K27ac) or repressive (H3K27me3) across subclasses. (N) Genome browser tracks of aggregated histone modifications, DNA methylation signals and chromatin states in PVALB and L6IT-1 subclass at example SLR loop anchors highlighted in (J), including loci with *TBR1, DLX1, SATB2*, and *RNF228*. (O) Heatmap showing GSEA of genes at SLR loop anchors using differential metrics between mature and progenitor cells from Keefe et al., 2025 as rank. Color indicates normalized enrichment score (NES) calculated from fgsea. Asterisk indicates significant association (Adjusted P value < 0.05). Box in all box plots shown this figure represents the interquartile range (IQR); whiskers extend to 2×IQR.

Consistent with the canonical role of enhancer-promoter communication, active loops significantly overlapped ABC-defined regulatory links (Figure S13B), and many of their target genes were associated with neuronal functions such as synaptic organization or membrane potential (Figure S13C). In contrast, repressive loops exhibited distinct organization: they spanned significantly longer genomic distances, contacted promoters less frequently (39.81% versus 58.81%), and preferentially connected transcriptionally silent loci, including genes involved in epithelial or kidney morphogenesis (Figure 6C, 6E and S13C). Notably, both active and repressive loops were enriched at domain-scale regulatory features, including super-enhancers and broad repressive domains (Figure 6F), suggesting that loop structure is coupled to higher-order regulatory features. Despite their lower abundance, repressive loops showed stronger interaction enrichment over local background than active loops (Figure 6G and 6H), consistent with more persistent, stabilized contacts.

Lineage-specific folding was especially evident at loci encoding key developmental regulators. For example, *DLX1* engaged contacts with multiple enhancers in PVALB interneurons (Figure 6I), where the gene is active. In contrast, these contacts were absent in glutamatergic neurons such as L6IT-1, where *DLX1* is silenced. Instead, this locus was embedded within a broad H3K27me3-marked domain spanning ∼300 kb (Figure 6I). Supporting a broader role for Polycomb-associated topology in constraining lineage programs, gene ontology analysis of repressive loop anchors in GABAergic neurons was enriched for genes implicated in alternative fate specification, including *SLC25A12* and *ITGA6* (Figure S13D)^108,109^. Together, these observations support a model in which H3K27me3-associated chromatin conformations sequester inappropriate regulatory programs through repressive 3D contacts.

While most loop analyses focus on interactions within local genomic neighborhoods, we identified a prominent class of “super long-range” (SLR) loops spanning >5 Mb (Figure 6J). These structures resemble Polycomb-associated long-range interactions reported in bulk neuronal Hi-C^110,111^, but single-cell resolution enabled deconvolution of their cell-type specificity. Using a supervised learning approach optimized to detect super long-range contacts (STAR Methods)^112^, we identified 12,130 SLR loops across 30 subclasses. SLR loops were highly cell-type specific, showing minimal overlap between neuronal subclasses such as PVALB and L6IT-1 (Figure 6K), and were preferentially detected in post-mitotic neurons while largely absent from astrocytes, oligodendrocytes, and microglia (Figure 6L).

To systematically characterize these structures, we annotated SLR loops by the epigenetic state of their anchors. Unlike canonical short-range loops, neuronal SLR loops were predominantly repressive, exceeding active SLR loops (44.87% vs 17.69%; Figure 6M). Genes engaged in SLR loops within a given subclass were significantly downregulated relative to subclasses without SLR loops, as illustrated by the reciprocal silencing of *TBR1, SATB2*, and *DLX1* between PVALB and L6IT-1 neurons (Figure 6N)^113-115^. Notably, SLR-associated targets were enriched for genes with strong activity in progenitor states during human cortical development (Figure 6O, S13D and S13E)^116^. These findings suggest that SLR interactions may encode a durable “developmental memory” by consolidating early developmental programs that must remain stably silenced in mature neurons. We therefore propose that, in post-mitotic neurons, subset of developmentally active genes are preferentially sequestered into SLR silencing loops, potentially stabilized within repressive nuclear environments, to ensure robust long-term repression of early-stage regulatory programs.

### Evolutionary Conservation and Divergence of Active and Repressive Regulatory Grammar

Comparative studies indicate that the cellular taxonomy of mammalian brain is broadly conserved across species, whereas species-specific traits likely arise largely from divergence in the noncoding regulatory genome^8,117-119^. Deciphering this *cis*-regulatory grammar is therefore essential for understanding the evolutionary origins of human cognition and for defining the translational limits of murine models.

To distinguish conserved epigenetic programs from human-biased specializations, we integrated the human data with matched Droplet Paired-Tag datasets from mouse cortex. Using a unified set of orthologous genes, we co-embedded transcriptomes across species and annotated 24 shared subclasses (Figure 7A and 7B). This analysis confirmed broad conservation of major neuronal and non-neuronal identities, with a notable exception of visual cortex-enriched L4IT neurons, consistent with the expansion and specialization of primate visual cortex relative to mouse.

**Figure 7.**
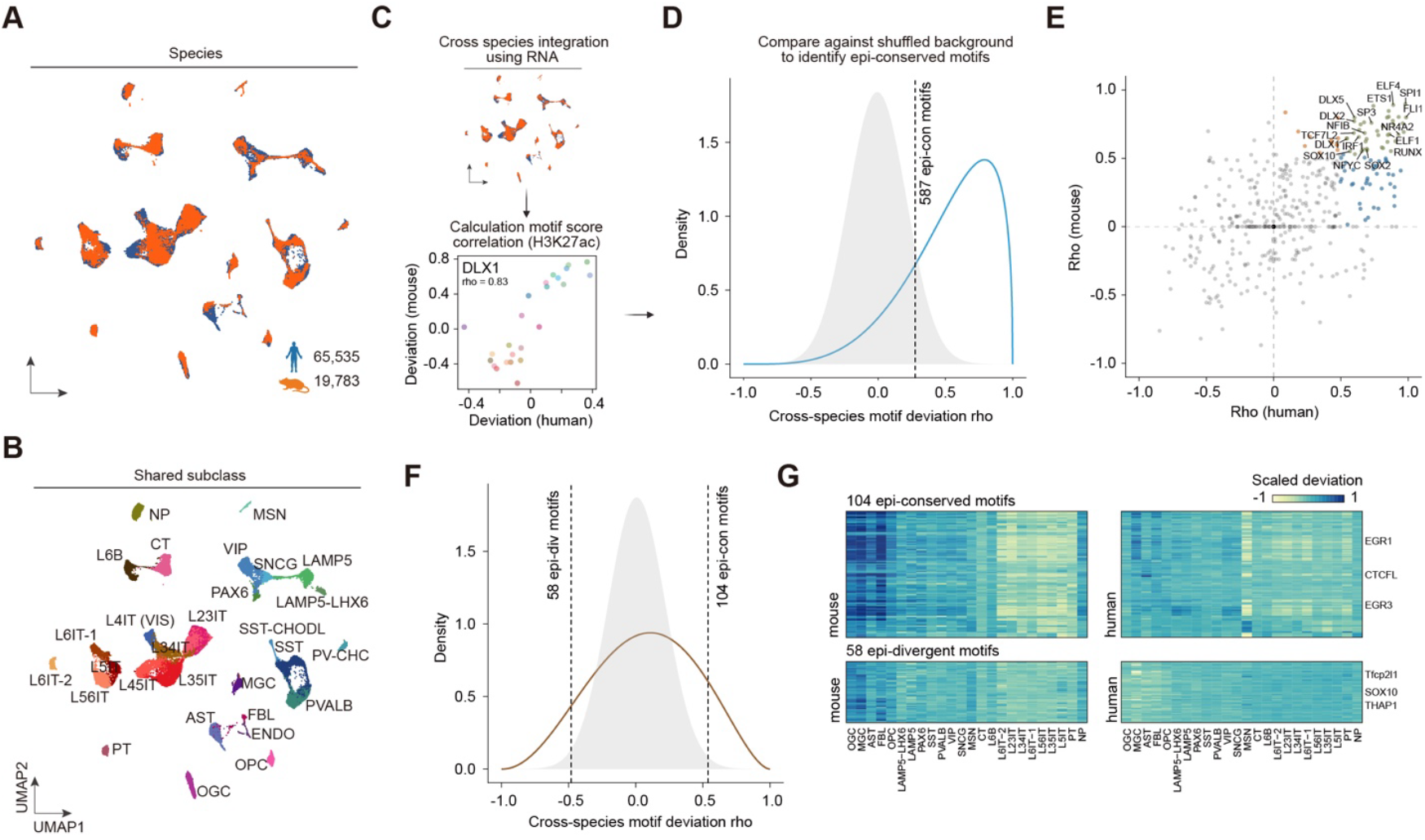
Evolutionary conservation and divergence of the regulatory grammar. (A-B) UMAP visualization of the integration of Droplet Paired-Tag transcriptomic data in cortical samples between human and mouse. Nuclei are colored by shared cell subclasses (A) or by species (B). (C) Schematic illustrating the process of identifying epi-conserved or divergent motifs across species. Following integration and shared subclasses annotation, the motif deviation score calculated using histone modification signals were averaged within each species across subclasses. Next, correlation between motif scores were calculated (e.g. DLX1) and rho cutoff was selected by comparing correlation distribution against shuffled background. Motifs with correlation larger than the cutoff were determined as epi-conserved motifs. (D) Distribution of cross-species motif correlation calculated with H3K27ac signal. Dashed line represents cutoff (FDR = 0.1) used to define epi-conserved motifs. (E) Scatterplot showing the correlation rho between motif score and expression of the associated transcription factor expression in human and mouse, for 587 H3K27ac-defined epi-conserved motifs. Strong correlation indicates potential gene regulatory programs. Top regulatory programs are labeled and colored based on whether they are specific to human (blue), mouse (orange), or both species (green). (F) Motif correlation density similar as (E) but calculated with H3K27me3 signal. Dashed lines represent cutoffs used to define epi-divergent (rho cutoff < 0) or epi-conserved (rho cutoff > 0) motifs. (G) Heatmap showing averaged motif deviation score across 104 H3K27me3-defined epi-conserved and 58 epi-divergent motifs in mouse (top) or human (bottom). Schematic of chromatin loop classification into Active, Repressive, and Super Long-Range (SLR) interactions.

To assess conservation of regulatory elements, we identified orthologous cCREs using reciprocal liftOver and quantified cross-species similarity of histone modifications within matched subclasses using generalized least squares (GLS) regression (STAR Methods)^119,120^. Across orthologous cCREs, divergence in histone modification signals was inversely related to the conservation index (Figure S14A and S14B). Comparison with chromatin accessibility further showed that ATAC and H3K27ac conservation scores were positively correlated but subject to constraints by distinct element type. Promoter-proximal cCREs were substantially more conserved than distal cCREs for chromatin accessibility, whereas this promoter-centric disparity was less pronounced for H3K27ac (Figure S14C and S14D). By contrast, H3K27me3 conservation showed no promoter-biased pattern, indicating distinct evolutionary constraints on activation and Polycomb-mediated repression.

We next examined conservation of transcription factor (TF) regulatory grammar. We developed a quantitative framework to classify “epi-conserved” and “epi-divergent” TF motifs by correlating motif deviation scores between human and mouse across matched subclasses (Figure 7C; STAR Method). Within the active chromatin landscape, motif activity was highly conserved (Figure 7D). We identified 587 epi-conserved motifs with comparable subclass-specific activity across species, including SOX5, ELF1, and NFIB in glia cells; ZBTB18, ESRRA, and NHLH1 in GABAergic neurons; and NEUROD1 in glutamatergic neurons (Figure S14E)^68^. To link motif activity to regulatory mechanisms, we compared motif deviation scores with RNA expression of the corresponding TFs and observed positive concordance for a substantial fraction of factors in both species (43 factors; 45.26%), including core lineage regulators such as DLX1, DLX5, NEUROD2, and TCF12 (Figure 7E)^121,122^. In contrast, the repressive regulatory grammar showed markedly greater evolutionary plasticity. Although 104 repressive motifs were identified as epi-conserved, 58 motifs were classified as epi-divergent which showed negative correlations between species (Figure 7F). Several transcription factors displayed human-specific or mouse-specific repressive motif activity, indicating rewiring of Polycomb-associated regulatory logic. One prominent example is THAP1, which is associated with hindbrain identity in humans but exhibits divergent repressive motif activity in mice (Figure 7G)^123,124^.

Together, these cross-species analyses reveal an evolutionary asymmetry in brain regulatory architecture. Active programs that establish core cellular identity are strongly conserved in mammals and are governed by a constrained TF grammar. In contrast, repressive programs exhibit substantially greater divergence, suggesting that species-specific evolution more often proceeds through rewiring of silencing mechanisms than through changes to core activation circuits. This divergence in repressive grammar may provide a flexible substrate for species-specific adaptations in brain development, plasticity and disease susceptibility.

## DISCUSSION

A central challenge in human brain genomics is to move beyond catalogs of molecular features and define how gene regulatory programs constrain cellular identity, plasticity, and disease vulnerability. By jointly profiling transcriptome with active and repressive histone modifications at single-cell resolution, this study provides a functional reference for the regulatory architecture of the adult human brain. Integrating chromatin state with three-dimensional genome organization, spatial information, and genetic variants further reveals organizing principles that transcriptomes or chromatin accessibility alone cannot resolve.

A key insight from this work is that regulatory state in the adult brain reflects coordinated deployment of activation and repression. H3K27ac profiles improves enhancer-gene linking, strengthens inference of active gene regulatory networks, and increases interpretability of disease-associated variants. In parallel, H3K27me3 signifies much more than inactivity. It reflects active suppression of alternative fates and earlier developmental programs. Crucially, repressive chromatin can preserve lineage history and prior signaling exposure after transcription stabilizes. This principle is most evident in oligodendrocytes, where histone landscapes reveal strong regional heterogeneity despite modest transcriptomic differences, and in SST interneurons where H3K27me3 gradients recapitulate gene programs consistent with distinct developmental and migratory histories. The coupling between repression and three-dimensional genome folding further indicates that this memory is not only biochemical but also architectural: neuron-enriched super long-range repressive loops preferentially engage genes that were active in progenitors but become durably silenced in post-mitotic neurons.

State-resolved annotation of *cis*-regulatory elements demonstrates that accessibility alone substantially overestimates regulatory potential. Only a subset of accessible elements are acetylated, evolutionarily constrained, and enriched for disease-associated variants. Conversely, Polycomb-repressed elements frequently overlap repetitive sequences and motifs for developmental transcription factors, consistent with a role in stabilizing mature cellular states by suppressing latent programs. Notably, cCREs that switch between active and repressed states across cell types form a particularly enriched regulatory core, suggesting that epigenetic variability, rather than constitutive activity, marks regulatory sequences most critical for specifying cell identity.

Integration of chromatin state with three-dimensional genome architecture clarifies how regulatory logic is embedded in nuclear topology. Active loops primarily support short-range enhancer-promoter communication, whereas repressive loops span longer distances and preferentially connect silenced loci. Most strikingly, neurons exhibit super long-range Polycomb-associated interactions that engage genes active in progenitor states but permanently silenced in mature lineages. These structures support a model in which developmental memory is consolidated not only through local biochemical mechanisms like repressive histone modifications or DNA methylation, but also through large-scale physical organizations like loops or hubs that insulates silenced programs against inappropriate reactivation.

Furthermore, spatial integration with MERFISH data extends these principles into tissue space. Laminar organization of the cortex coincides with graded transitions in chromatin state, including coordinated shifts in activation and repression that mirror or even precede transcriptional changes. These findings indicate that chromatin state encodes positional and historical information that contributes to regional specialization while maintaining overall cellular identity in the adult brain.

These findings motivate a predictive view of chromatin state. Regulatory state is not simply a snapshot of the ongoing activity. It can determine future responsiveness by delimiting which programs remain inducible and which are insulated by durable repression. This provides hypotheses for why certain central nervous system cell types retain selective plasticity despite overall stability, and why stress, aging, or disease may preferentially destabilize specific gene modules in a cell-type- and region-restricted patterns^1,125-128^.

The regulatory framework established here also advances the interpretation of noncoding genetic variants. By anchoring genetic risk to cell-type-specific active regulatory elements, enhancer-gene links, and GRNs, this resource moves beyond proximity-based heuristics toward connecting variants to effector genes and vulnerable cell types. By partitioning heritability across cell-type-specific active elements, intersecting fine-mapped variants with enhancer-gene links, and prioritizing allelic effects using a sequence-based model of epigenetic impact, this study provides an integrated framework from statistical associations to candidate regulatory elements, effector genes, implicated cell types, and putative transcription factor. More broadly, combining these regulatory maps with patient-specific genetics or perturbation assays should accelerate the translation from variants to mechanisms, particularly for developmental and neurodegenerative disorders in which spatiotemporal specificity underlies pathology and therapeutic opportunity.

Finally, comparative analysis between human and mouse reveals a striking evolutionary asymmetry. Active regulatory programs remain strongly conserved across mammals, consistent with their roles in specifying broad cell identity. In contrast, repressive grammar is more plastic, suggesting that species-specific adaptations may preferentially emerge through rewiring of silencing programs. This asymmetry may provide explanation why mouse models reproduce many core neuronal identities yet incompletely capture human-biased constraints on developmental programs, plasticity, and disease susceptibility.

In summary, joint single-cell profiling of transcription and chromatin state enables functional resolution of the human brain regulatory genome. By linking activation, repression, 3D topology, spatial organization, and genetic variants within a unified framework, this resource provides a roadmap for decoding how epigenetic regulation shapes brain function, evolution, and disease.

### Limitations of the study

Several limitations warrant consideration. The present study focuses on adult postmortem tissue from a limited number of male donors and histone marks, and future work incorporating additional epigenetic modifications, developmental stages, perturbations, and direct spatial epigenomic assays will be essential to refine chromatin-state models. Nevertheless, the integrative framework presented here establishes a foundation for understanding how regulatory state governs both stability and adaptability in the human brain.

## ACKNOWLEDGMENTS

This publication was supported by and coordinated through the Brain Initiative Cell Atlas Network (BICAN) and funded by the National Institute of Mental Health (UM1MH130994). We thank E. Melief, A. Schantz, K. Kern, A. Keen, and L. Keene at the University of Washington BioRepository and Integrated Neuropathology (BRaIN) Laboratory for outstanding technical and administrative support with tissue collection, preservation, and characterization. We are deeply grateful to the brain donors and their families, without whom this work would not be possible. This publication was also supported by the Flow Cytometry Core Facility of the Salk Institute (RRID:SCR 014839) with funding from NIH-NCI CCSG: P30 CA01495, and Shared Instrumentation Grants S10-OD023689 (Aria Fusion cell sorter) and S10 OD034268 (Thermo Fisher Bigfoot). This publication also includes data generated at the UC San Diego IGM Genomics Center utilizing an Illumina NovaSeq X Plus that was purchased with funding from a National Institutes of Health SIG grant (#S10 OD026929). We also thank the Broad Institute sequencing cores for generating sequencing data and the QB3 MacroLab at UC Berkeley for purification of Tn5 transposase and recombinant RNase inhibitor. Z.W. is a D.D.Brown Awardee of the Life Sciences Research Foundation. We also thank Yang Eric Li, Jingtian Zhou, and Xiaotao Wang for valuable discussions and suggestions.

## AUTHOR CONTRIBUTIONS

Study conception: Y.X., B.R.

Study supervision: M.M.B., B.R.

Brain collection: R.D.H., C.D.K., X.X.

Tissue dissection and sample preparation: J.A.R., T.B., A.S.B., J.K.W., K.W.K., J.L., S.C., K.G.R., C.K.Y., J.A., Y.S., A.B., N.S., C.B., C.C., W.O., C.O., M.L., M.V.M., C.R., S.N.A., N.E., J.O., M.M.B.

Droplet Paired-Tag data generation: L.C., A.L., H.S.I., K.D., T.L., Y.X., Z.W.

MERFISH data generation: J.F., C.T., A.M., S.M., Q.Z.

Sequencing data analysis: Y.X., G.Z., W.D., E.A., K.L., B.H., D.L.

Imaging data analysis: Y.X., E.B., C.K., Z.Z., A.K., J.O., Q.Z.

Data interpretation: Y.X., L.C., G.Z., T.W., J.R.E., M.M.B., B.R.

Manuscript writing: Y.X., L.C., M.M.B., B.R.

All authors edit and approve the final manuscript.

## DECLARATION OF GENERATIVE AI AND AI-ASSISTED TECHNOLOGIES

During the preparation of this manuscript, the authors used OpenAI’s GPT-5.2 model for language editing and proofreading to improve clarity and readability. The authors reviewed and revised all output as needed and took full responsibility for the content of the publication.

