## Supplementary Figures for "Single-Cell Atlas of Transcription and Chromatin States Reveals Regulatory Programs in the Human Brain"

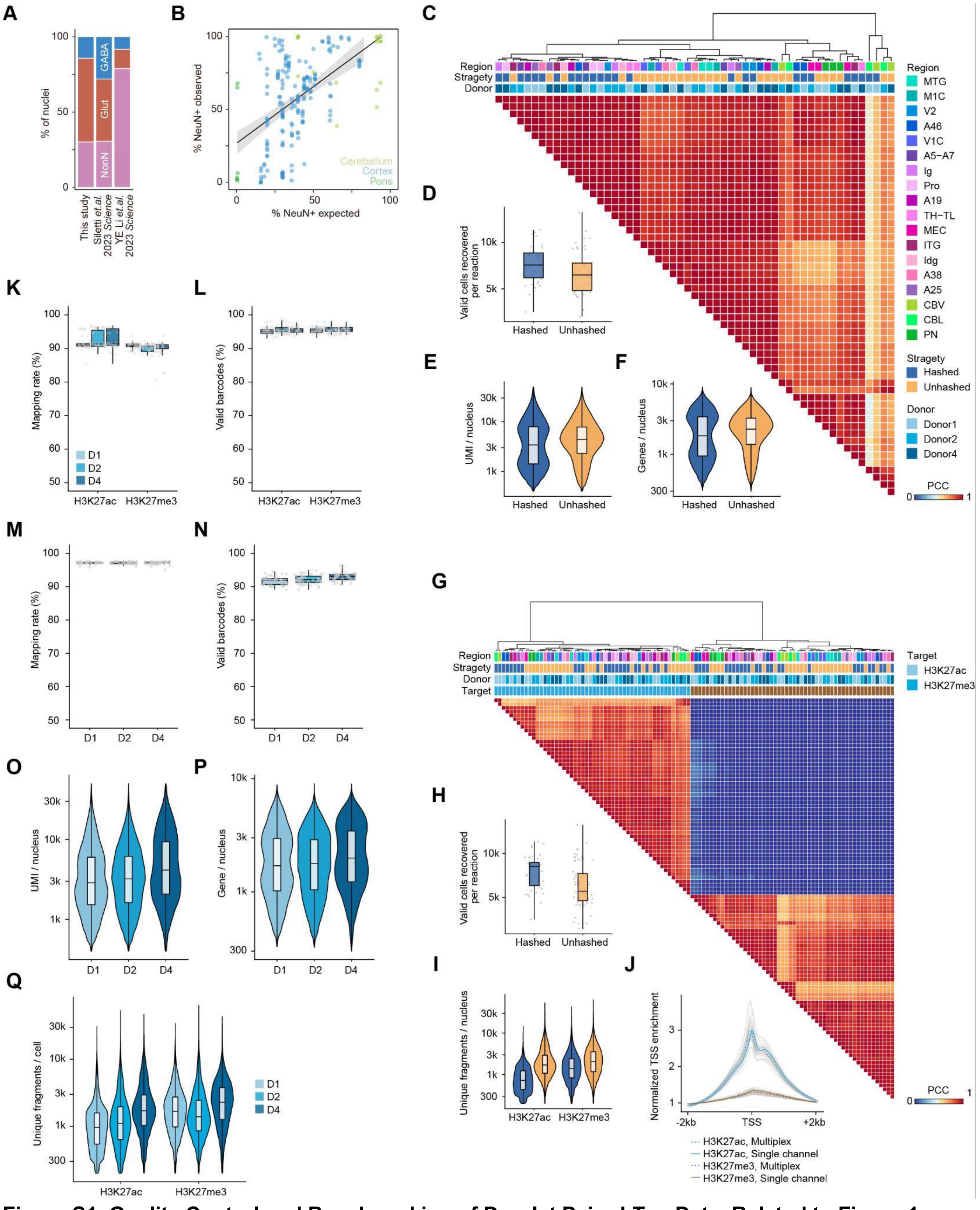

**Figure S1. Quality Control and Benchmarking of Droplet Paired-Tag Data, Related to Figure 1**
(A) Comparison of cell class composition (Glutamatergic, GABAergic, Non-neuronal) obtained in this study versus published human brain atlases with (Siletti *et al.*, 2023) or without (Li *et al.*, 2023) NeuN enrichment.

(B) Scatter plot showing the correlation between observed and expected NeuN+ nuclei percentages per sample.
(C) Pearson correlation coefficient matrix of transcriptome profiles pseudobulked by experimental conditions, showing mixing of donors and processing strategies (i.e., hashed vs. unhashed).
(D-F) Box plots displaying quality control metrics for the transcriptome (RNA) modality, comparing hashed and unhashed samples the number of valid nuclei recovered (D), Unique Molecular Identifiers (UMI) per nucleus (E), and genes detected per nucleus (F).
(G) Pearson correlation coefficient matrix of histone modification profiles similar to (C), showing mixing of donors and processing strategies but clear separation of histone marks profiled. Dots in (D) represent the number of nuclei recovered per library.
(H-I) Box plots displaying quality control metrics for the DNA modality (H3K27ac and H3K27me3), showing the number of valid nuclei recovered (H) and unique fragments per nucleus (I). Dots represent the number of nuclei recovered per library.
(J) Normalized Transcription Start Site (TSS) enrichment scores for H3K27ac and H3K27me3 libraries, stratified by multiplexing strategy. Each line represents an individual library. Colored lines represent normalized TSS calculated by pooling all sequencing reads in each condition.
(K-L) Box plots showing mapping rates per library (K) and percentage of valid barcodes per library (L) for histone modification libraries across donors. Each dot represents a sequencing library.
(M-N) Mapping rates (M) and percentage of valid barcodes (N) per library for RNA libraries, similar to K-L.
(O-Q) Violin plots and box plots showing the distribution of data quality on the single cell level, including UMIs per nucleus (O), genes per nucleus (P), and unique fragments per cell (Q) stratified by donor or histone modifications. Violin shape represents kernel density estimation. For all box plots in the figure, the center line represents the median, box limits represent the upper and lower quartiles (75th and 25th percentiles), and whiskers represent 2× interquartile range (IQR).

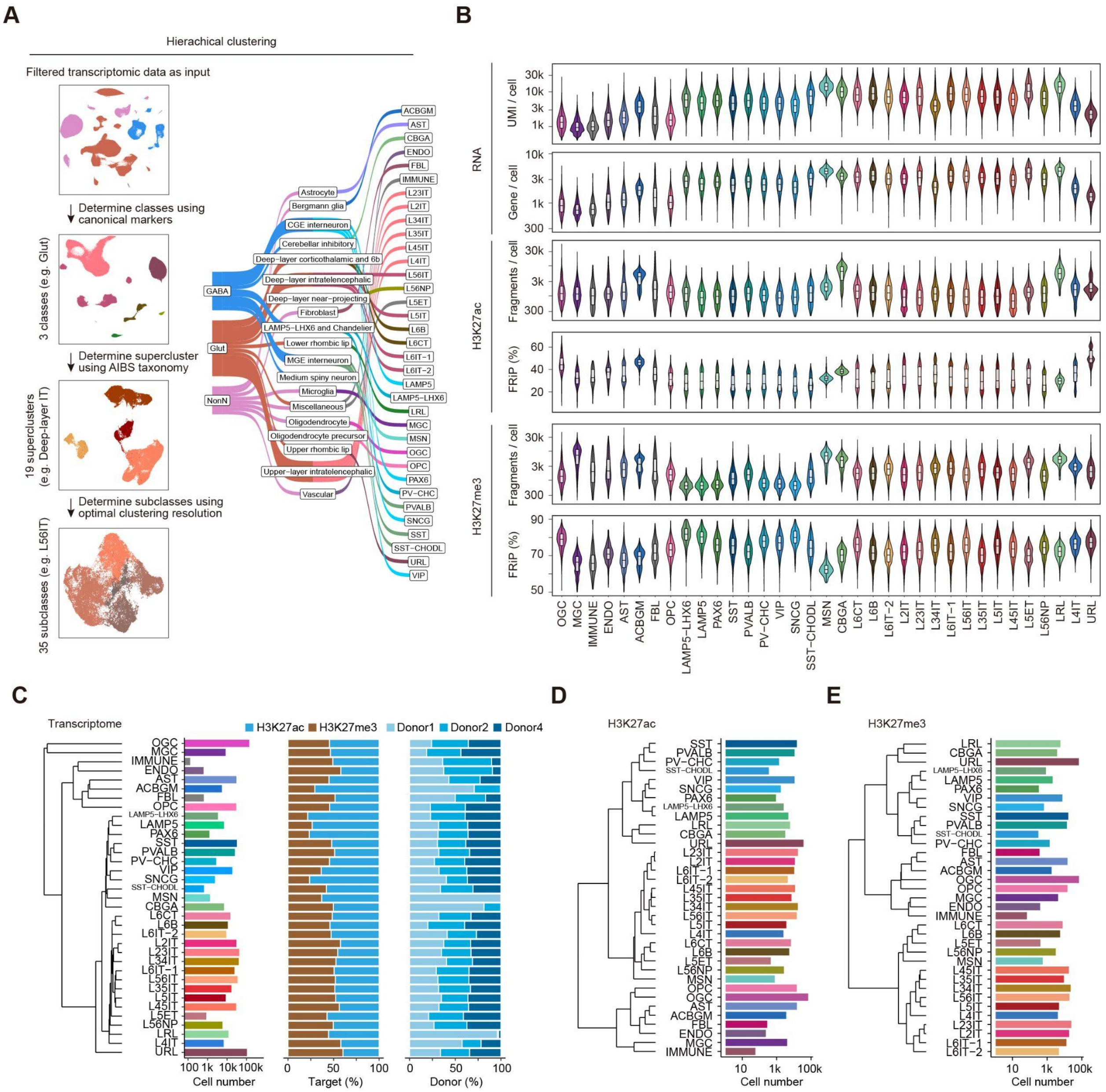


**Figure S2. Cell Type Taxonomy and Subclass-Specific Quality Metrics, Related to Figure 1**.

(A) Diagram showing the workflow of hierarchical clustering and dendrogram of the 35 annotated subclasses, grouped into 19 superclusters and 3 major classes.

(B) Violin plots and boxplots displaying quality control metrics including UMI / nucleus, genes / nucleus, unique fragments / nucleus, and Fraction of Reads in Peaks (FRiP) / nucleus for RNA, H3K27ac, and H3K27me3 modalities across all annotated subclasses. For all box plots, the center line represents the median, box limits represent the upper and lower quartiles (75th and 25th percentiles), and whiskers represent 2× interquartile range (IQR).

(C) Left: Hierarchical clustering of transcriptome profiles pseudobulked by cell subclasses. Middle: Bar plot showing the total number of nuclei recovered for each subclass. Right: Bar plots showing the proportion of nuclei by histone modification targeted or by donor.

(D) Left: Hierarchical clustering of H3K27ac profiles pseudobulked by cell subclasses. Right: Bar plots showing the total number of H3K27ac nuclei recovered for each subclass.

(E) Similar plot as (D) but for H3K27me3.

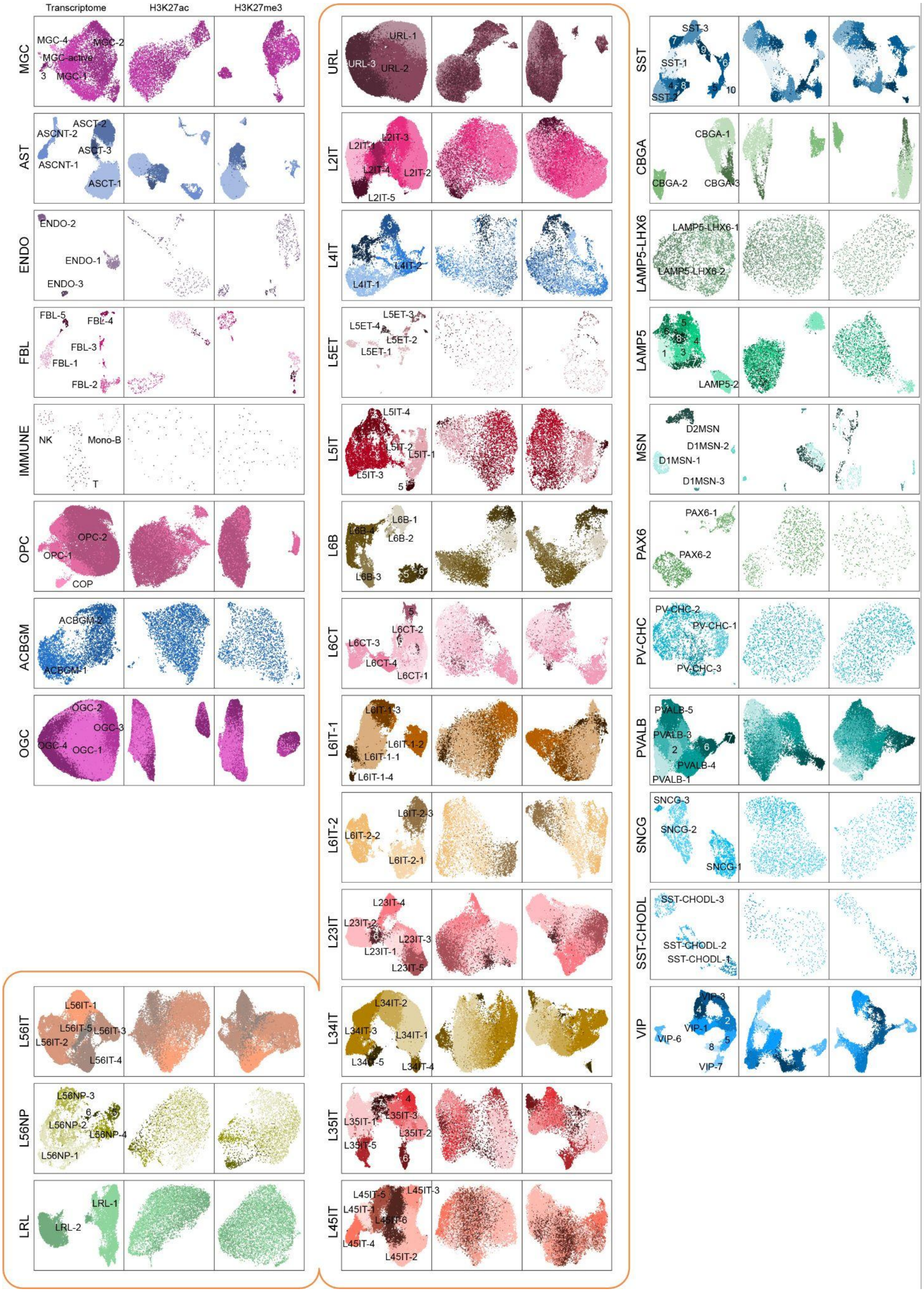
Transcriptome
H3K27ac
H3K27me3
MGC
MGC-4
MGC-2
MGC-active
3
MGC-1
AST
ASCT-2
ASCNT-2
ASCT-3
ASCNT-1
ASCT-1
ENDO
ENDO-2
ENDO-1
ENDO-3
FBL
FBL-5
FBL-4
FBL-3
FBL-1
FBL-2
IMMUNE
NK
Mono-B
T
OPC
OPC-2
OPC-1
COP
ACBGM
ACBGM-2
ACBGM-1
OGC
OGC-2
OGC-3
OGC-4
OGC-1
L56IT
L56IT-1
L56IT-5
L56IT-3
L56IT-2
L56IT-4
L56NP
L56NP-3
L56NP-2
L56NP-4
L56NP-1
LRL
LRL-1
LRL-2
URL
URL-1
URL-3
URL-2
L2IT
L2IT-3
L2IT-1
L2IT-4
L2IT-2
L2IT-5
L4IT
L4IT-2
L4IT-1
L5ET
L5ET-3
L5ET-4
L5ET-2
L5ET-1
L5IT
L5IT-4
L5IT-2
L5IT-1
L5IT-3
L6B
L6B-1
L6B-4
L6B-2
L6B-3
L6CT
L6CT-2
L6CT-3
L6CT-4
L6CT-1
L6IT-1
L6IT-1-3
L6IT-1-2
L6IT-1-1
L6IT-1-4
L6IT-2
L6IT-2-3
L6IT-2-2
L6IT-2-1
L23IT
L23IT-4
L23IT-2
L23IT-3
L23IT-1
L23IT-5
L34IT
L34IT-2
L34IT-3
L34IT-1
L34IT-5
L34IT-4
L35IT
L35IT-3
L35IT-1
L35IT-2
L35IT-5
L45IT
L45IT-5
L45IT-3
L45IT-1
L45IT-6
L45IT-4
L45IT-2
SST
SST-3
SST-1
SST-2
CBGA
CBGA-1
CBGA-2
CBGA-3
LAMP5-LHX6
LAMP5-LHX6-1
LAMP5-LHX6-2
LAMP5
LAMP5-2
MSN
D2MSN
D1MSN-2
D1MSN-1
D1MSN-3
PAX6
PAX6-1
PAX6-2
PV-CHC
PV-CHC-2
PV-CHC-1
PV-CHC-3
PVALB
PVALB-5
PVALB-3
PVALB-4
PVALB-1
SNCG
SNCG-3
SNCG-2
SNCG-1
SST-CHODL
SST-CHODL-3
SST-CHODL-2
SST-CHODL-1
VIP
VIP-3
VIP-1
VIP-6
VIP-7

**Figure S3. UAMP visualization for cell clusters for each subclass, Related to Figure 1**.
Uniform Manifold Approximation and Projection (UMAP) embeddings of cell subclasses using H3K27ac, and H3K27me3 modalities, each further divided into cell clusters with colors indicating cluster identity.

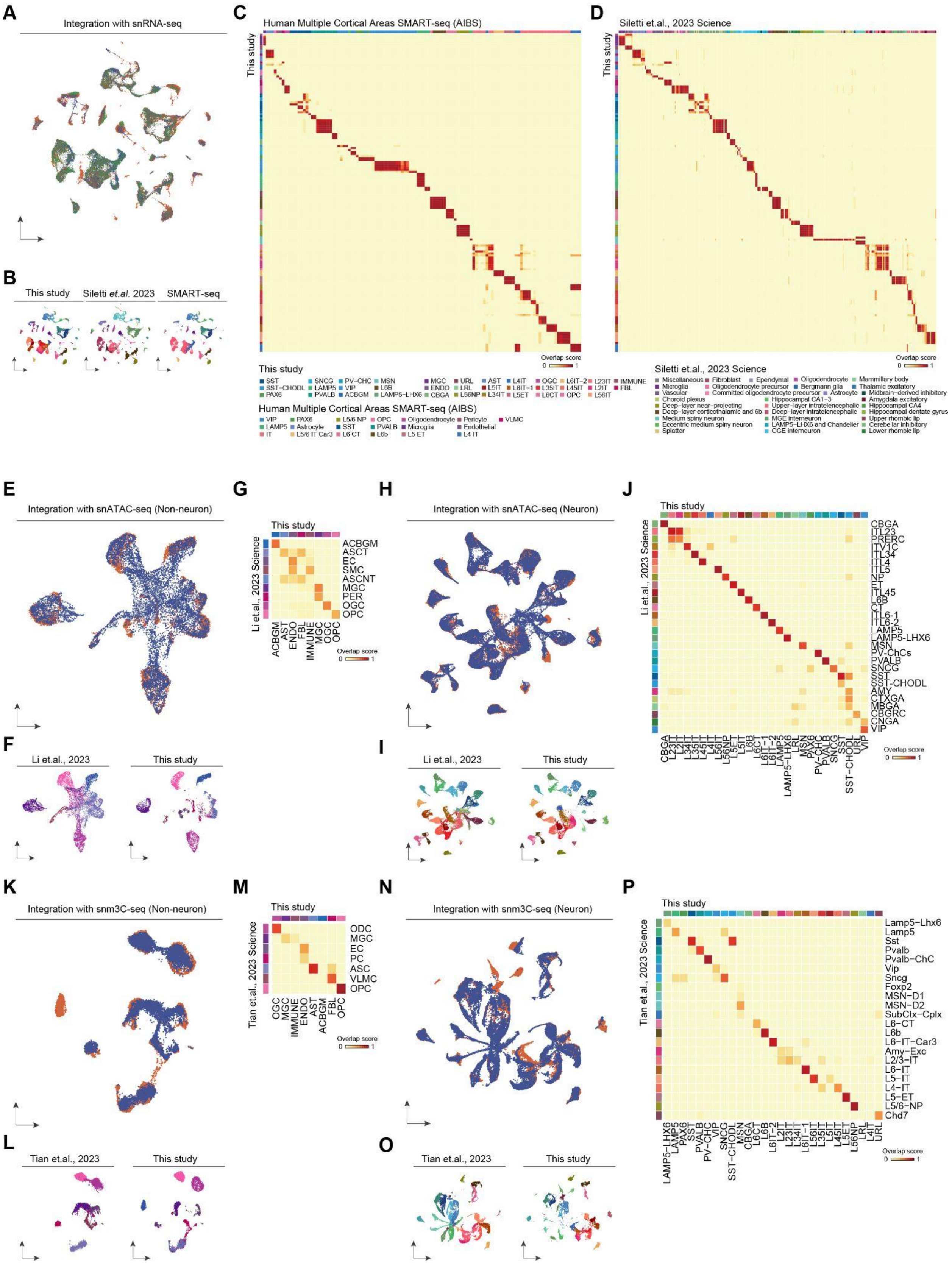


**Figure S4. Multi-Omic Integration with Published Human Brain Atlases, Related to Figure 1**

(A) UMAP visualization of the integration between this study's snRNA-seq data and external datasets, including whole human brain snRNA-seq (Siletti *et al.*, 2023) and human multiple cortical areas SMART-seq (AIBS SMART-seq).
(B) UMAP visualization of the subclass annotation for each dataset on the integrative embedding.
(C-D) Confusion matrix showing the overlap score between cell cluster annotations in this study versus the AIBS SMART-seq atlas (C) or Siletti *et al.,* 2023 (D).
(E) UMAP visualization of the integration results between non-neuronal cells in this study and snATAC-seq data from Li *et al.,* 2023.
(F) UMAP visualization of the subclass annotation of non-neuronal cells for each dataset on the integrative embedding.
(G) Confusion matrix showing the overlap score between non-neuronal subclass annotations in this study versus Li *et al.*, 2023.
(H-I) UMAP visualizations similar to (E) and (F) but shown for neuronal cells.
(J) Confusion matrix similar as (G) but for neuronal subclasses.
(K-L) UMAP visualization of the integration result and subclass annotations for non-neuronal cells in this study and in snm3C-seq data from Tian *et al.,* 2023.
(M) Overlap score heatmap for non-neuronal annotations in this study versus Tian *et al.,* 2023.
(N-O) UMAP visualizations similar to (K) and (L) but shown for neuronal cells
(P) Overlap score heatmap for neuronal cell annotations in this study versus Tian *et al.,* 2023.

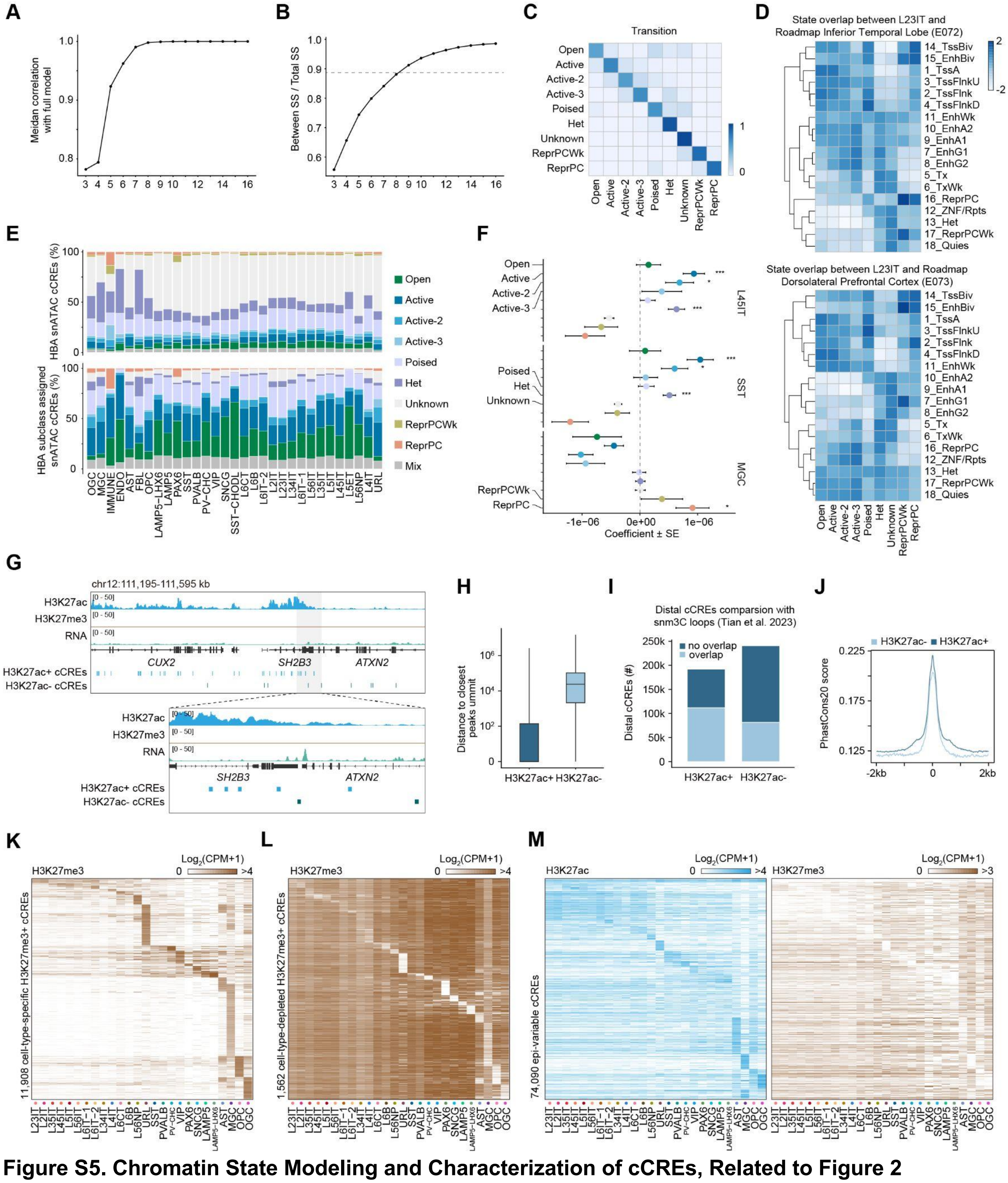


**Figure S5. Chromatin State Modeling and Characterization of cCREs, Related to Figure 2**

(A) Line plots showing the median Pearson correlation coefficients (PCC) of the states in all models with the full model (16-state).

(B) K-means classification performed on the emission probabilities from all models. The optimal number of states was defined as the smallest k at which cluster separation reached at least 95% (grey dashed line) of the maximum separation observed across all k values. Cluster separation was assessed using the sum of squares (SS).

(C) Heatmap showing the transition probability matrix between nine chromatin states.

(D) Heatmap showing the genomic overlap (log2 observed/expected) between chromatin states annotated in L23IT neurons and reference human brain tissues annotated with the 18-state models from the Epigenome Roadmap. Enrichment of genomic overlap is scaled by row.
(E) Stacked bar plot showing the proportion of cCREs from Li *et al.* 2023 assigned to each chromatin state in this study. Results for all cCREs (top) as well as cCREs accessible in each cell subclass (bottom) are shown.
(F) Coefficients from LDSC analysis quantifying the association of Schizophrenia trait to cCREs annotated in each chromatin state for select cell types (MGC, SST, L45IT). Error bars represent standard error (SE). Asterisks indicate significance calculated using Benjamini-Hochberg correction. ***, FDR < 0.0005; **, FDR < 0.005; *, FDR < 0.05.
(G) Example genome browser track at the *ATXN2* locus showing H3K27ac+ and H3K27ac- cCREs alongside histone modification and RNA tracks.
(H) Box plots showing the genomic distance (log scale) of H3K27ac+ and H3K27ac- cCREs to the closest H3K27ac peak summit.
(I) Bar plots comparing the number of distal cCREs (H3K27ac+ vs H3K27ac-) overlap with snm3C-seq loops identified in Tian *et al.* (2023).
(J) Line plots showing the distribution of PhastCons conservation scores centered on H3K27ac+ versus H3K27ac- cCREs.
(K) Heatmap of H3K27me3 signal (log2 CPM+1) for 11,908 cell-type-specific repressive cCREs.
(L) Heatmap of H3K27me3 signal (log2 CPM+1) for 562 cell-type-depleted repressive cCREs.
(M) Heatmap of H3K27ac (left) and H3K27me3 (right) signal (log2 CPM+1) for 74,090 epi-variable cCREs across subclasses. For all box plots in the figure, the center line represents the median, box limits represent the upper and lower quartiles (75th and 25th percentiles), and whiskers represent 2× interquartile range (IQR).

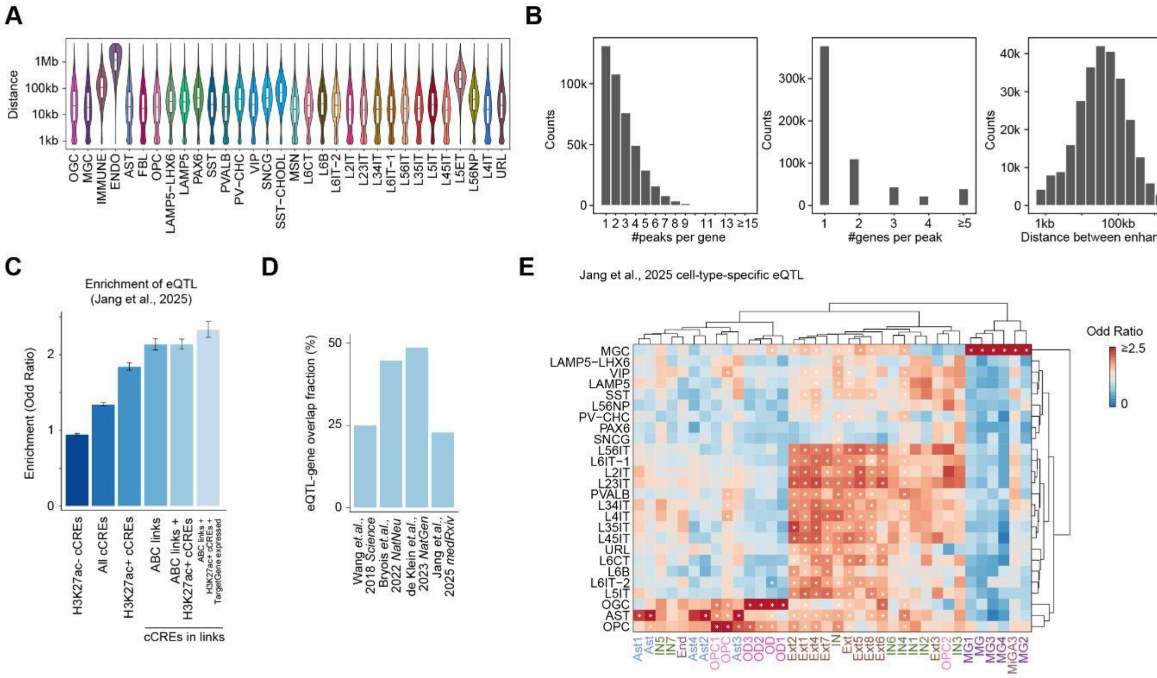


**Figure S6. Properties of Enhancer-Gene Links from ABC model, Related to Figure 3**

(A) Violin plots and box plots showing the distribution of distances between ABC-linked enhancers and transcription start sites (TSS) for all cell subclasses before filtering. The box center line represents the median, box limits represent the upper and lower quartiles (75th and 25th percentiles), and whiskers represent 2× interquartile range (IQR).
(B) Histograms showing the number of peaks per gene (left), genes per peak (middle), and the distribution of log scaled enhancer-TSS genomic distances (right) for filtered ABC links.
(C) Bar plots showing the enrichment (Odds Ratio) of eQTLs (Jang *et al.*, 2025) within different categories of cCREs. Error bars represent mean ± standard deviation (SD). Odds Ratio are calculated using Fisher's exact test.
(D) Bar plot comparing the fraction of eQTL-gene overlap in this study with other published eQTL datasets. Calculation is already based on shared eQTL (*i.e.,* eQTL located in enhancer regions).
(E) Heatmap of cell-type-specific enrichment of eQTLs-gene links (Jang *et al.*, 2025) within ABC inferred enhancer-gene links in each subclass. Color scale represents Odds Ratio. Asterisks indicate significance calculated by Fisher's exact test. and corrected using Benjamini-Hochberg (BH) method.

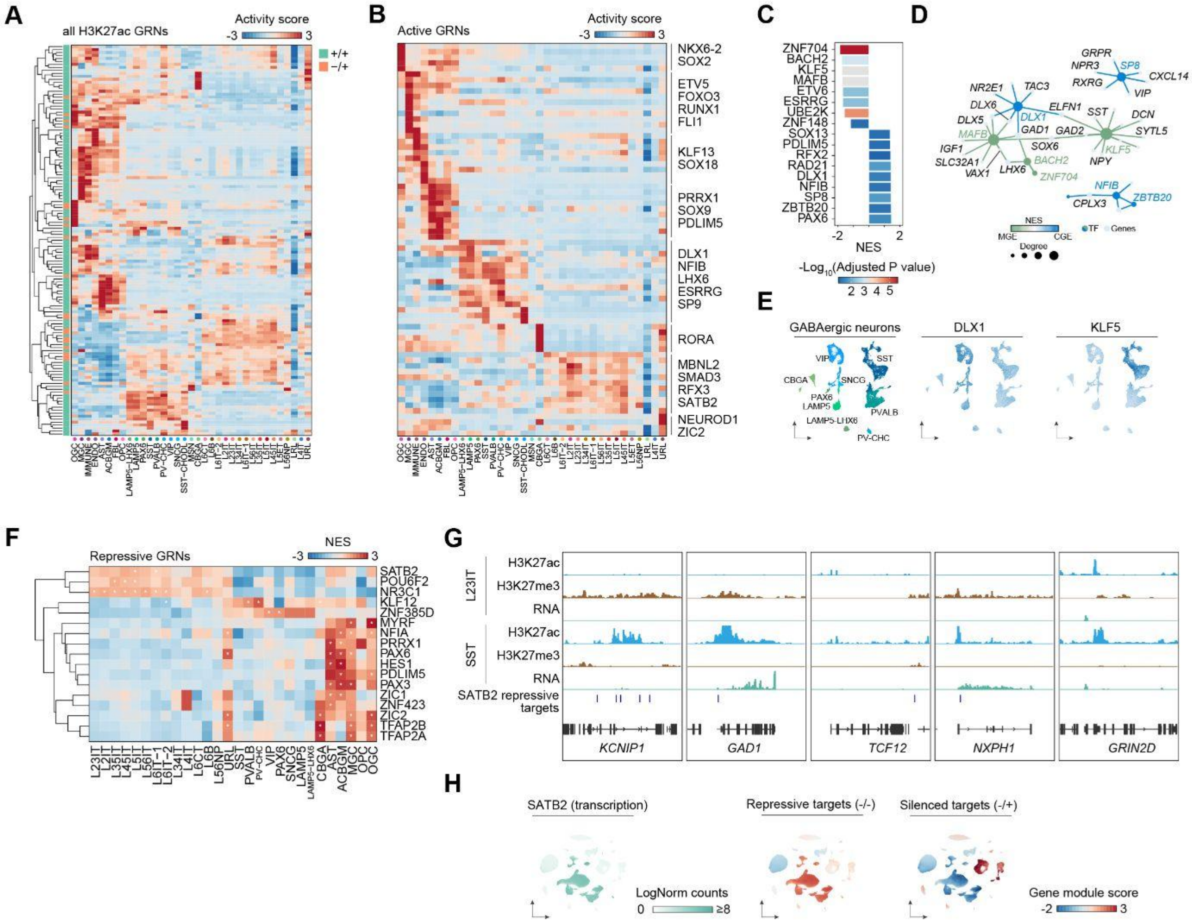


**Figure S7. Active and Repressive Gene Regulatory Network Analysis, Related to Figure 3**
(A) Heatmap of TF activity scores for all identified H3K27ac-associated gene regulatory networks (GRNs).

Colors indicate activity values computed for each eRegulon as √(enhancers H3K27ac signal × target genes transcription), and summarized across eRegulons within each GRN. Row annotations denote GRN type and whether TF activity is positively (+/+) or negatively (−/+) correlated with target gene expression.

(B) Heatmap of TF activity scores for active GRNs enriched in at least one cell subclass. Enrichment was assessed by GSEA using differential expression statistics to rank all genes, with GRN target genes as the gene set. P values are calculated by fGSEA and corrected for multiple testing using the BH method.

(C) Bar plots showing the normalized enrichment score (NES) of MGE/CGE-associated TF GRNs, calculated using GSEA similar as described in (B). P values are calculated by fGSEA and adjusted using the BH method.

(D) Network visualization of representative MGE / CGE-associated TFs and their top targeting genes. Node size represents connectivity (Degree), node color represents identity (TF or genes). For TF nodes, color indicates whether the TF is associated more with MGE or CGE interneurons.

(E). UMAP visualization of GABAergic neurons clustering results, colored by cell subclass (left) or enrichment scores for selected GRNs (right).

(F) Heatmap of NES of repressive GRNs defined by H3K27me3 in each subclass, calculated using GSEA similar as described in (B). Asterisk indicates significance (FDR < 0.05). P values are calculated by fGSEA and corrected using the BH method.

(G) Genome browser tracks showing the histone modification and RNA signal in L23IT versus SST on example SATB2 repressive GRNs targeted genes. Regions associated with repressive GRNs were annotated alongside.

(H) Left: UMAP visualization of the *SATB2* transcription level across all nuclei. Right: UMAPs colored by the module scores calculated with genes in SATB2 repressive (-/-) or silenced (-/+) GRN.

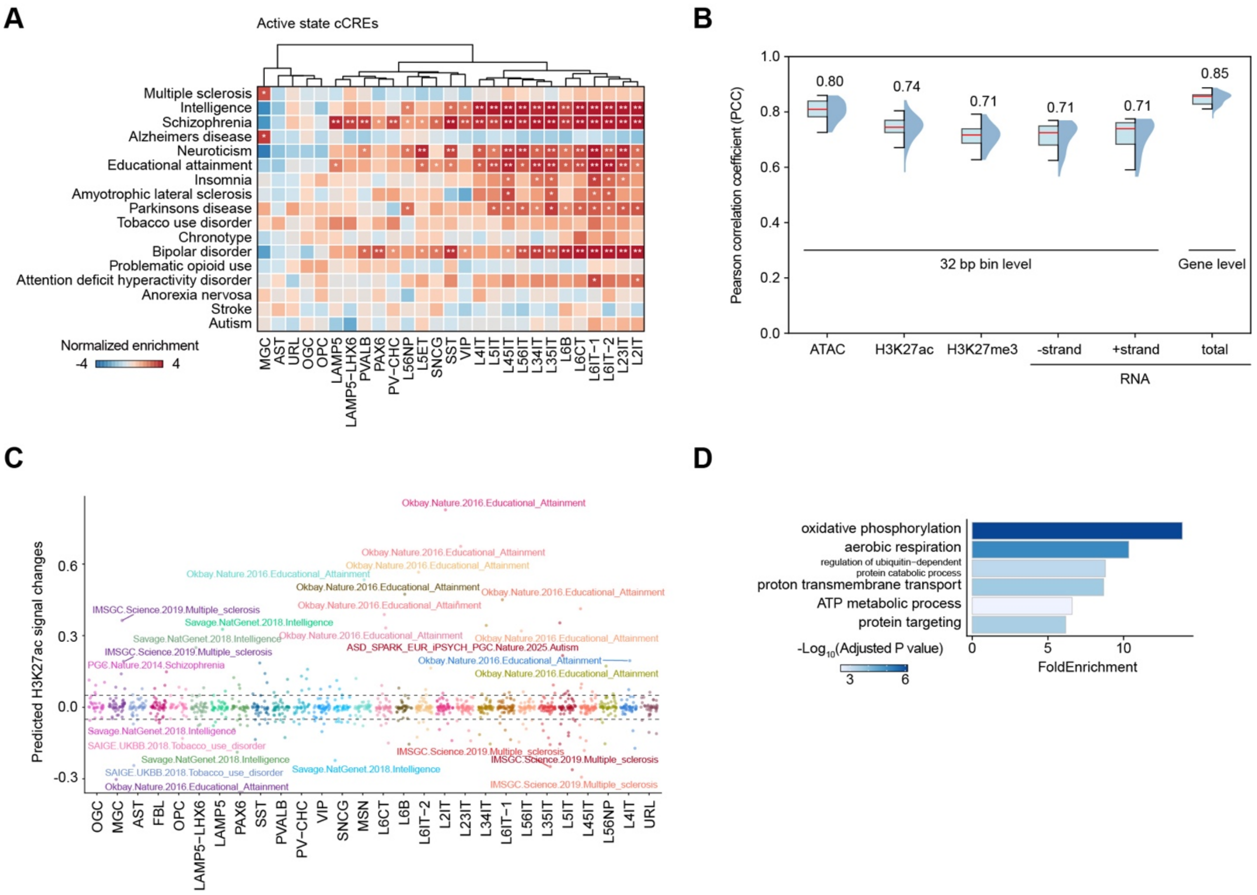

**Figure S8. Disease Heritability and Variant Effect Prediction, Related to Figure 3**

(A) Heatmap showing LDSC heritability enrichment for representative neuropsychiatric and neurological traits within active states cCREs across cell subclasses.Color indicates Z-score. P values are corrected to FDR using BH correction. Asterisk indicates significance (FDR < 0.05).

(B) Box plots and violin plots showing Pearson correlation coefficients (PCC) quantifying the model prediction accuracy with test dataset for histone modifications and RNA signal. Histone modification signals are aggregated by 32bp genomic bin for training and prediction, while total transcription is aggregated at gene levels. Box plot center line: median; box limits: IQR; whiskers: 1.5×IQR.

(C) Scatter plots showing the predicted H3K27ac signal changes over fine-mapped variants across cell subclasses. Example traits associated with strong effect variants are labeled.

(D) Gene Ontology enrichment analysis for genes associated with NFE2L1 GRN from SCENIC+ analysis. P-values are calculated using a hypergeometric test in enrichGO.

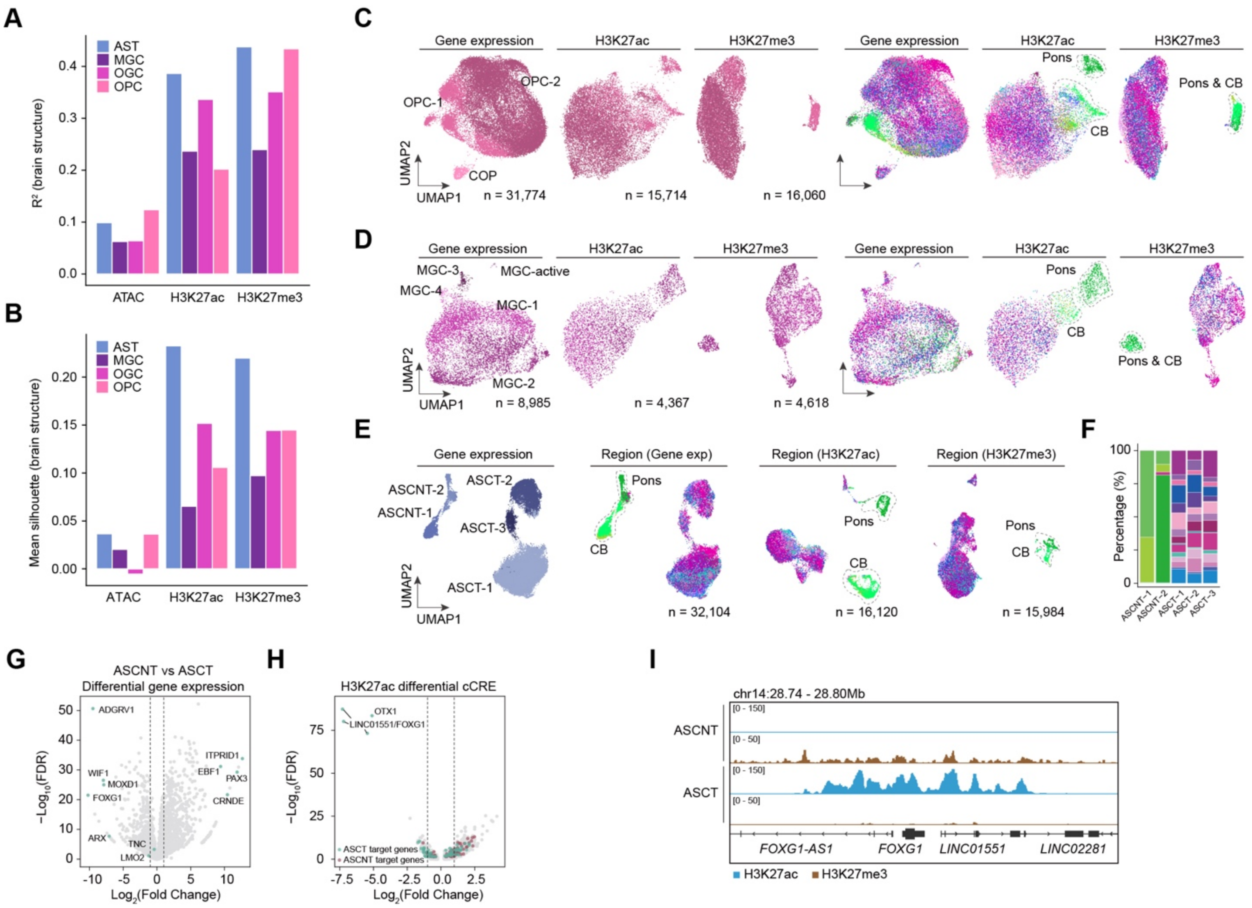


**Figure S9. Regional Heterogeneity in Glial Subclasses, Related to Figure 4**

(A) Bar plots showing the variance explained ($R^2$) by brain structure for astrocytes (AST), microglia (MGC), oligodendrocytes (OGC), and oligodendrocytes progenitors (OPC).

(B) Bar plots of the mean silhouette score for brain structure clustering across modalities.

(C-E) UMAP visualization of transcriptome, H3K27ac, and H3K27me3 profiles on OPC (C), MGC (D) and AST (E), colored by cell cluster (left) or brain region (right).

(E) UMAP visualization using transcriptome, H3K27ac, and H3K27me3 data on AST, colored by cell cluster annotation or brain region.

(F) Stacked bar plot showing the regional composition of AST cell clusters.

(G) Volcano plot of differential gene expression between cortical ASCT versus ASCNT. Canonical marker genes are colored and labeled. Dashed lines represent absolute log2 FoldChange = 1.

(H) Volcano plot of differential H3K27ac cCREs between ASCT and ASCNT. cCREs targeting genes that are shown to be differentially expressed (FDR < 0.01, absolute log2 FoldChange ≥ 1) between ASCT and ASCNT are colored in green or red, respectively. Targeting genes of top cCREs are labeled. Dashed lines represent absolute log2 FoldChange = 1.

(I) Genome browser track of the *FOXG1* locus in AST, showing specific H3K27ac signal in ASCT and repressive H3K27me3 in ASCNT.

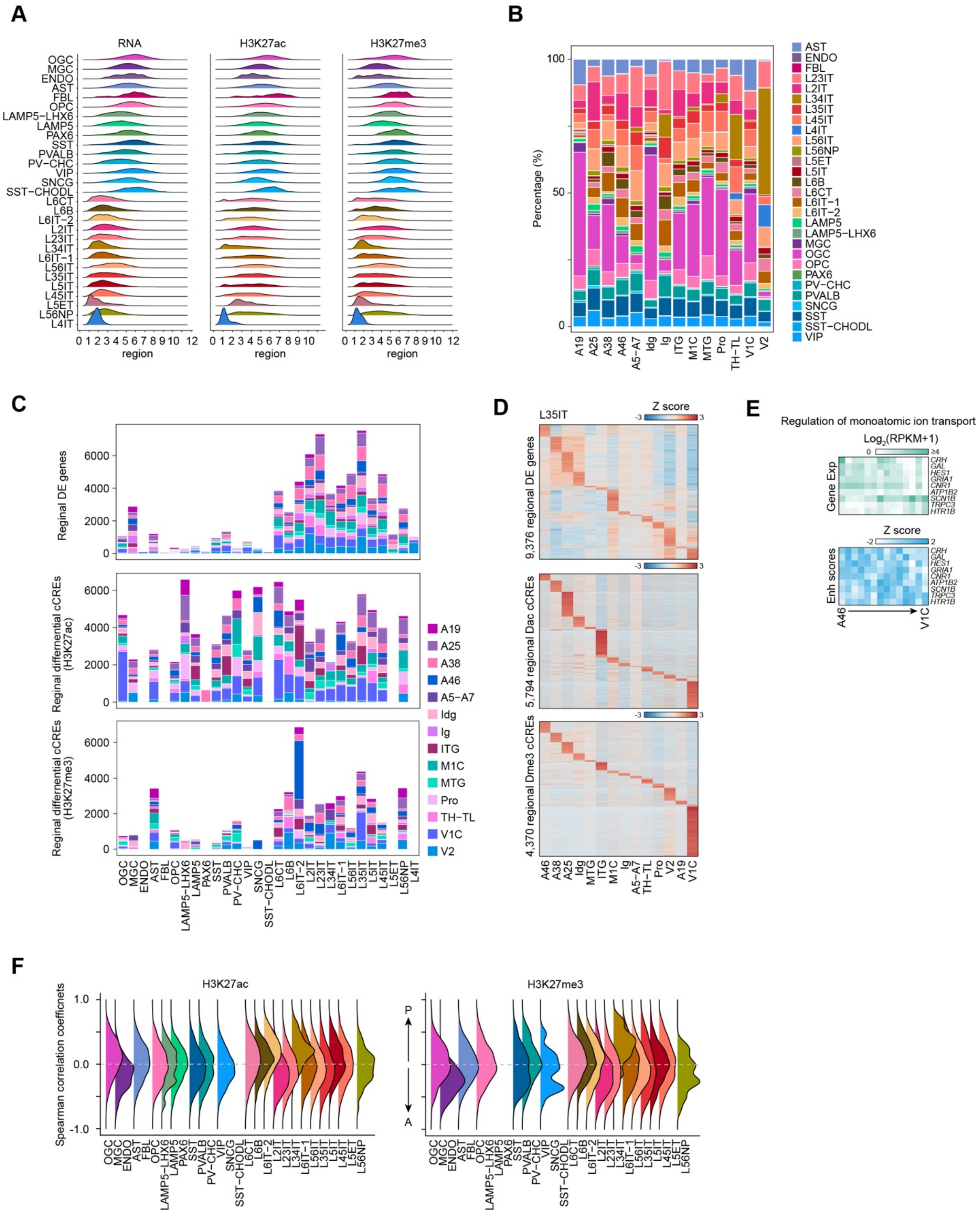


**Figure S10. Systematic Analysis of Regional Heterogeneity Across Cortex, Related to Figure 4**

(A) Ridge plots showing the LISI distribution of RNA, H3K27ac, and H3K27me3 signals across cortical regions for all subclasses.

(B) Stacked bar plots showing regional composition of all cell subclasses across cortical areas.

(C) Bar plots showing the number of regional differentially expressed genes (top), differential cCREs calculated using H3K27ac (middle) and H3K27me3 (bottom) for each subclass. Color indicates the source of regions.
(D) Heatmaps of transcription signal (top) for regional differential genes and histone modification signals (bottom) for differential cCREs in L35IT neurons, ordered by cortical region. Signals are Z-score normalized.
(E) Heatmaps showing aggregated transcription signal and enhancer scores for example genes in the GO term "Regulation of monoatomic ion transport" across regions.
(F) Ridge plots of Spearman's rank correlation coefficients quantifying the cCRE signal correlation with the anterior-posterior (A-P) axis across subclass. Subclasses with few (<10) or without regional differential cCREs are not displayed.

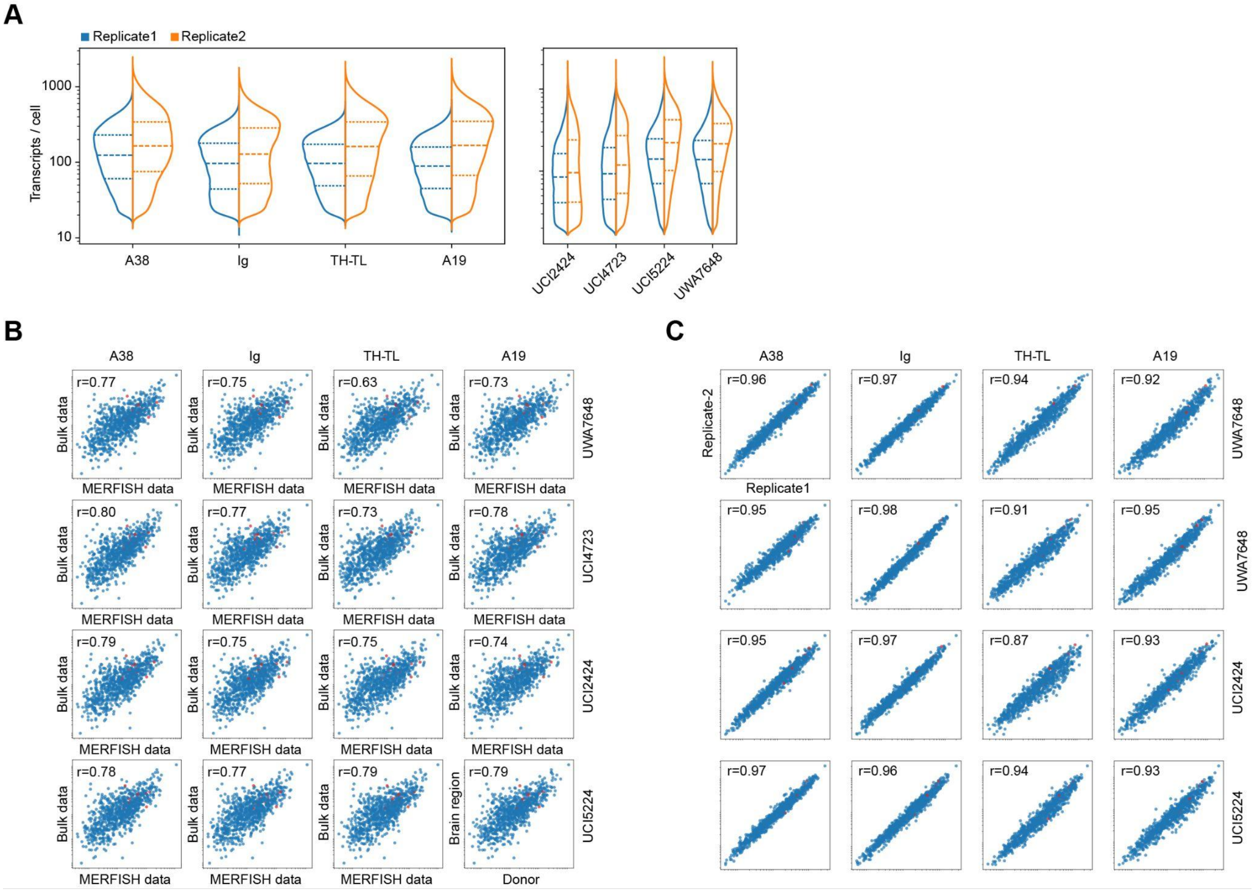


**Figure S11. Quality Control and Validation of MERFISH Spatial Transcriptomics Data, Related to Figure 5**
(A) Violin plots displaying the distribution of detected transcripts per cell across different brain regions (A38, Ig, TH-TL, A19) and donors (UCI2424, UCI4723, UCI5224, UWA7648) used for MERFISH experiments, stratified by biological replicates.
(B) Scatter plots showing the correlation between pseudo-bulk gene expression profiles derived from MERFISH (x-axis) and standard bulk RNA-seq data (y-axis) for matched samples. r denotes Pearson correlation coefficient.
(C) Scatter plots comparing average gene expression between biological replicates across regions and donors. r denotes Pearson correlation coefficient.

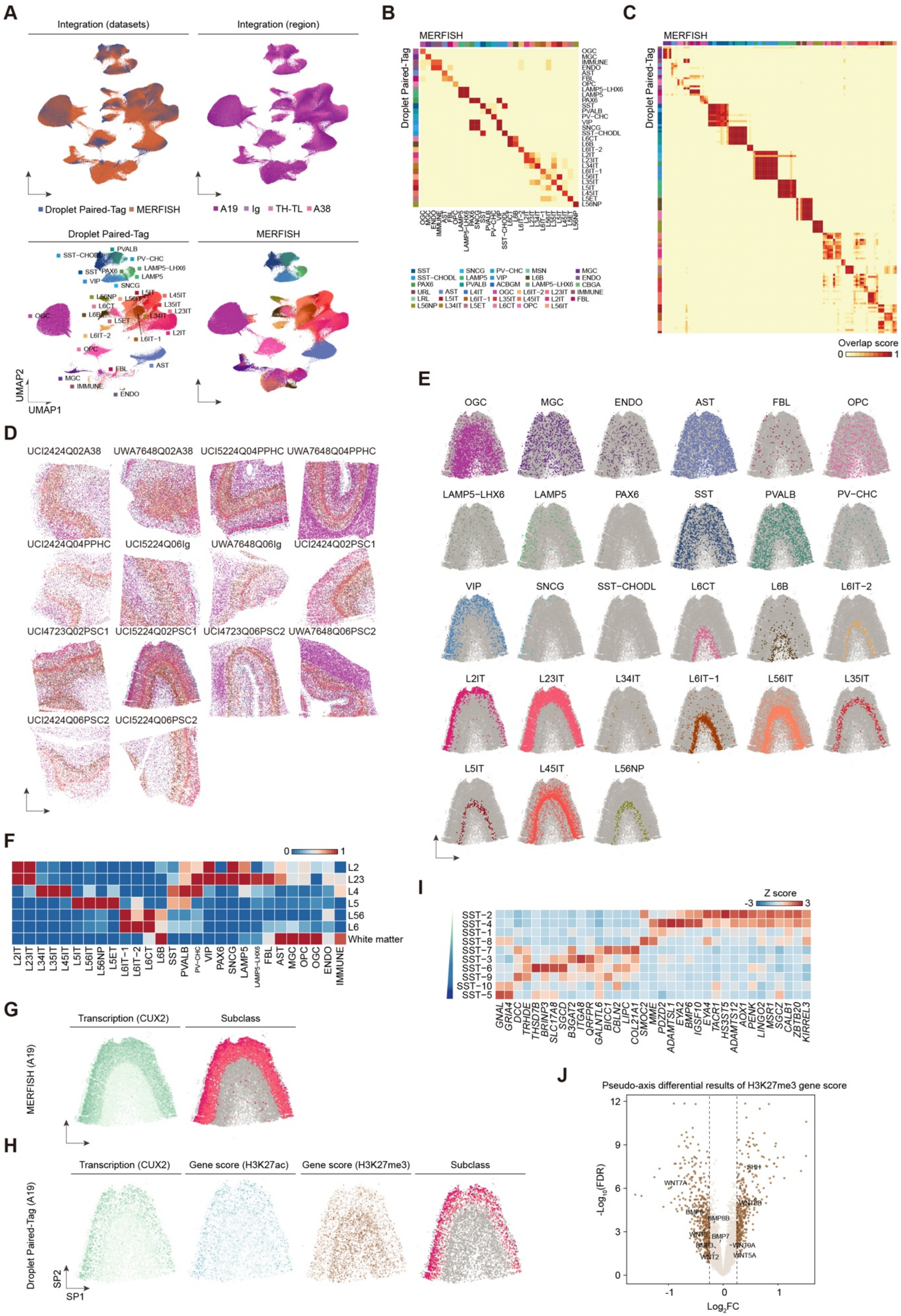
A
Integration (datasets)
Integration (region)
Droplet Paired-Tag
MERFISH
A19
Ig
TH-TL
A38
UMAP1
UMAP2
B
C
Droplet Paired-Tag
Overlap score
0
1
D
UCI2424Q02A38
UWA7648Q02A38
UCI5224Q04PPHC
UWA7648Q04PPHC
UCI2424Q04PPHC
UCI5224Q06Ig
UWA7648Q06Ig
UCI2424Q02PSC1
UCI4723Q02PSC1
UCI5224Q02PSC1
UCI4723Q06PSC2
UWA7648Q06PSC2
UCI2424Q06PSC2
UCI5224Q06PSC2
E
OGC
MGC
ENDO
AST
FBL
OPC
LAMP5−LHX6
LAMP5
PAX6
SST
PVALB
PV−CHC
VIP
SNCG
SST−CHODL
L6CT
L6B
L6IT−2
L2IT
L23IT
L34IT
L6IT−1
L56IT
L35IT
L5IT
L45IT
L56NP
F
L2
L23
L4
L5
L56
L6
White matter
I
Z score
SST−2
SST−4
SST−1
SST−8
SST−7
SST−3
SST−6
SST−9
SST−10
SST−5
G
Transcription (CUX2)
Subclass
MERFISH (A19)
H
Gene score (H3K27ac)
Gene score (H3K27me3)
Droplet Paired-Tag (A19)
SP1
SP2
J
Pseudo-axis differential results of H3K27me3 gene score
−Log10(FDR)
Log2FC
WNT7A
SHH
WNT8B
BMP6
BMP8B
BMP7
WNT9A
WNT5A
WNT2

**Figure S12. Spatial Integration Framework and Laminar Epigenetic Heterogeneity, Related to Figure 5**

(A) UMAP visualization of the integration results of Droplet Paired-Tag and MERFISH data using shared transcriptomic features. Nuclei are colored based on the datasets (top left), brain regions (top right) and annotated subclasses in each dataset (bottom).

(B-C) Confusion matrices displaying the overlap score between cell subclass (B) and cell cluster (C) annotations in the Droplet Paired-Tag and the MERFISH dataset.

(D) Spatial visualization of high quality MERFISH slides across different donors and regions. Cells are colored based on the predicted cell subclass.

(E) Spatial plots of individual cells in each annotated subclass on representative tissue sections. Distinct laminar layering is visible especially for glutamatergic neuron subclasses.

(F) Heatmap showing the normalized enrichment of major predicted cell subclasses within cortical layers (L2 to L6 and White Matter) defined by MERFISH spatial coordinates.

(G) Spatial visualization of layer-specific marker *CUX2* for L2IT and L23IT neurons on MERFISH data.

(H) Spatial visualization of *CUX2* transcription, H3K27ac gene score, and H3K27me3 gene score on Droplet Paired-Tag nuclei with imputed spatial position.

(I) Heatmap of transcription signals for example pseudodepth-associated marker genes for 10 identified SST cell clusters. Signals are Z score normalized.

(J) Volcano plot showing differential H3K27me3 gene scores along the SST pseudo-depth trajectory. Positive fold change indicates enrichment of repression in superficial layers. Developmental genes including Wnt/Bmp pathway members (*WNT7A*, *BMP7*, *SHH*) are labeled. Dashed lines represent absolute log2 FoldChange = 0.25.

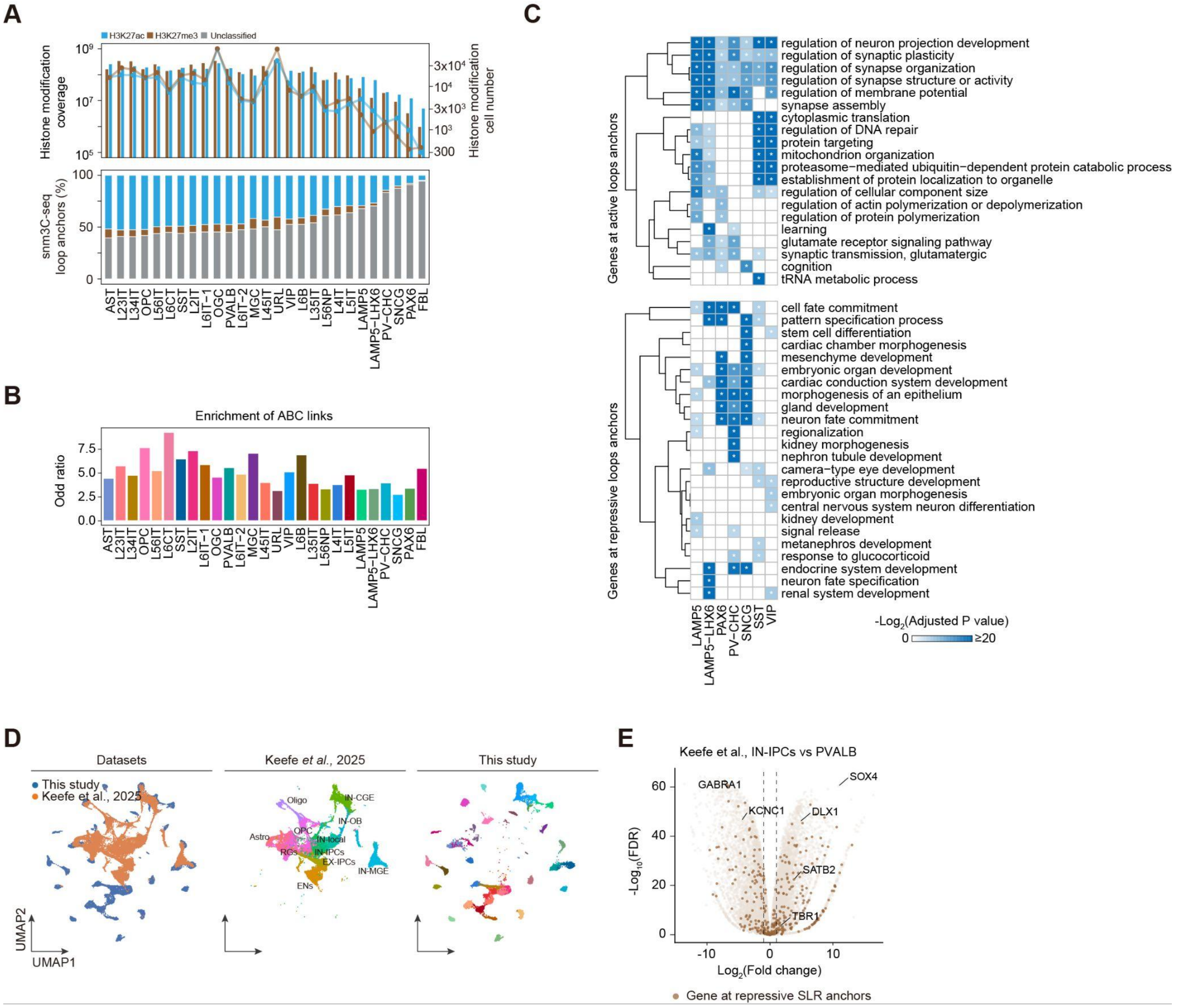


**Figure S13. Functional Characterization of Active and Repressive Chromatin Loops, Related to Figure 6**

(A) Top: Bar plots showing the length of histone modification narrow peaks (coverage) and the nuclei number in each histone modification modality. Bottom: Proportion of loop anchors stratified by their histone modification status (H3K27ac, H3K27me3, or unclassified).

(B) Bar plot showing the overlap enrichment (Odd ratio) of active loops with ABC links. P-value calculated by Fisher's exact test.

(C) Gene Ontology analysis for genes engaged in active chromatin loops (top) or repressive loops (bottom) across GABAergic neuron subclasses. P-values are calculated using a hypergeometric test in enrichGO and adjusted for multiple testing using BH correction. Asterisk indicated significance (adjusted P value < 0.05).

(D) UMAP visualization of the integration results between this study and developmental human brain data (Keefe *et al.*, 2025). Nuclei are colored based on datasets (left), cluster annotation in Keefe *et al.* (middle), and subclass annotation in this study (right).

(E) Volcano plot showing differential gene expression between mature PVALB interneurons (this study) and developmental inhibitory neuron progenitors (Keefe *et al.*, 2025). Representative genes engaged in Super Long-Range (SLR) repressive loops in mature cells are labeled. Dashed lines represent absolute log2 FoldChange = 1.

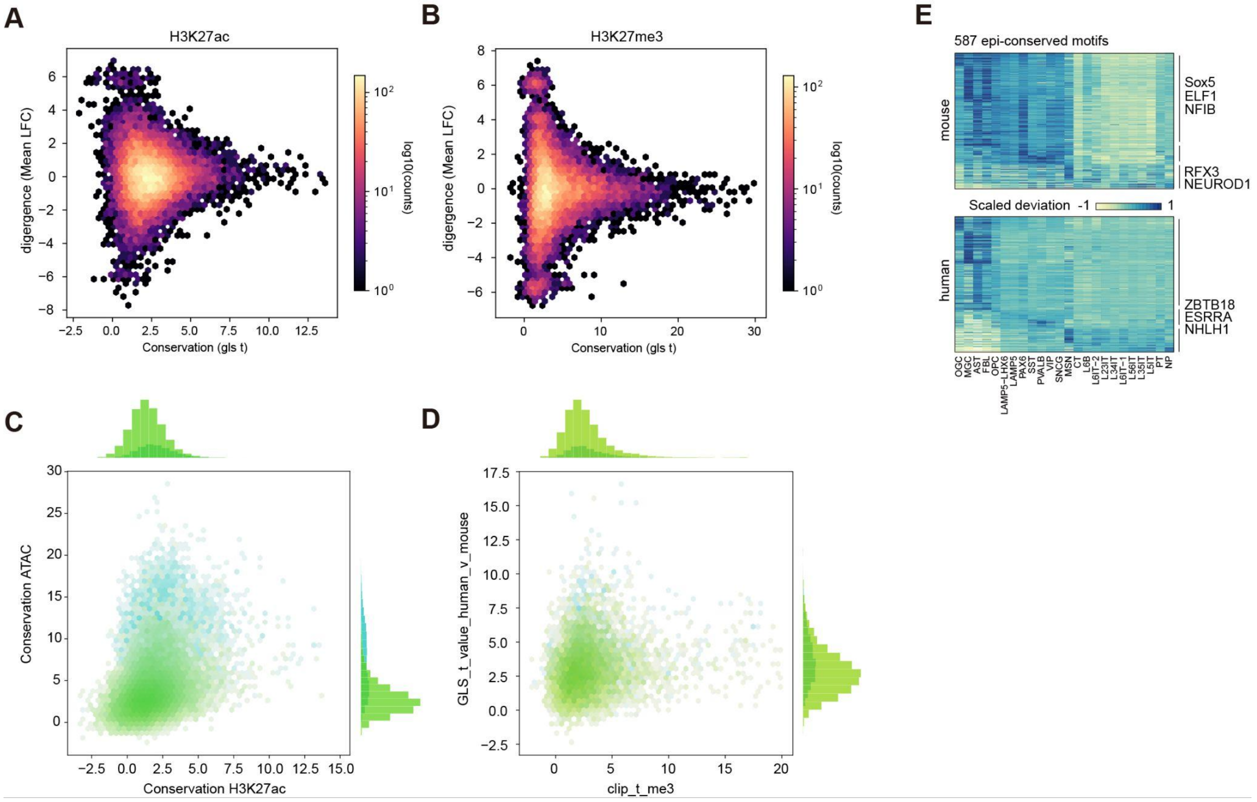


**Figure S14. Cross-Species Epigenomic Conservation Metrics, Related to Figure 7**

(A-B) Scatter plots comparing the epigenetic conservation score (GLS t-statistic, x-axis) versus transcriptional divergence between species (Mean Log Fold Change, y-axis) for H3K27ac (A) and H3K27me3 (B) cCREs. Brighter colors indicate higher density of cCREs.

(C) Scatter plot showing the positive correlation between conservation scores calculated using H3K27ac versus ATAC-seq data. Histogram summarizing the density distribution of conservation score for each modality. Colors represent different groups of genomic features (blue, promoter-proximal cCREs; green, distal cCREs).

(D) Scatter plot showing the modest correlation between conservation scores calculated using H3K27me3 (clip_t_me3) versus ATAC-seq data (GLS_t_value_human_v_mouse) similar as shown in (C).

(E) Paired heatmaps showing the scaled motif deviation scores for 587 epi-conserved motifs in mouse (top) and human (bottom). Rows represent motifs and columns represent matched cell subclasses. Representative motifs in different cell classes are displayed. Color indicates scaled motif deviation score.
